# A widespread internal brain state for fentanyl withdrawal

**DOI:** 10.64898/2026.05.04.722791

**Authors:** Karim Abdelaal, Kathryn K. Walder-Christensen, Cameron Blount, Kellie M. Williford, Marisella Adams-Grimaldi, Stephen D. Mague, David E. Carlson, Kafui Dzirasa

## Abstract

Opioid addiction is characterized by escalating drug use, driven in part by negative reinforcement from withdrawal, but the neural processes linking withdrawal to increased drug-taking remain poorly understood. Here, we use multisite local field potential recordings and interpretable machine learning to identify large-scale brain networks engaged by repeated opioid exposure and withdrawal. After discovering that repeated fentanyl exposure induces a network of widespread high beta and low gamma oscillations, we then identified a distinct brain network that selectively encodes the emergence and severity of opioid withdrawal. This latter network, termed *EN-Withdrawal*, is characterized by regional gamma oscillations and widely synchronized delta/theta oscillations. Its activity patterns predict the emergence of spontaneous and naloxone-precipitated withdrawal across multiple independent cohorts, generalizing across mice, sex, opioids, and dosing regimens, while persisting over multiple days of withdrawal. Using a novel, data-driven severity index, we find that network activity scales with individual behavioral severity without simply reflecting ongoing somatic behaviors or general aversion, suggesting that *EN-Withdrawal* underlies a withdrawal-induced internal state. Strikingly, network activity predicts the escalation of fentanyl self-administration on a mouse-by-mouse basis in experienced, but not drug-naïve, animals. These findings reveal a neurophysiological substrate of the negative reinforcement cycle of addiction that shapes individual vulnerability.

## INTRODUCTION

Opioid use disorder (OUD) remains one of the most pressing public health crises of our time, contributing to over 74,000 overdose deaths and an estimated $2.7 trillion economic burden in the United States in 2023 alone^1^. Opioid dependence develops following repeated opioid exposure and is characterized by physiological adaptations that alter affective^2–4^, motivational^5–7^, and cognitive processes^8–10^, ultimately driving tolerance and escalation of drug use through a recurring process termed the addiction cycle^11–13^. Throughout the progression from casual use to compulsive opioid abuse within this cycle, withdrawal avoidance plays a critical role in driving sustained drug seeking and relapse through a negative reinforcement process^14,15^. Individuals with OUD experience a constellation of physical and psychological symptoms during opioid withdrawal, which are consistently identified as a primary barrier to recovery^16,17^. Given the widespread expression of opioid receptors^18^, this constellation of symptoms likely reflects disruptions of large-scale neural dynamics. However, the neural underpinnings by which withdrawal influences the trajectory of opioid abuse remain poorly understood.

The neural mechanisms underlying opioid withdrawal have been studied extensively at molecular, cellular, and circuit levels^3,11,12,14,15,19–23^. Recruitment of stress-related systems for negative emotional regulation within the extended amygdala^24–26^, including corticotrophin-releasing factor^4,19^ and dynorphin^27–30^ signaling, contribute to the negative affective components of withdrawal, while hyperactivity of noradrenergic locus coeruleus ensembles drive many somatic opioid withdrawal symptoms^18,31–36^. More recently, distinct cellular populations within the central amygdala have been shown to respond to naloxone-precipitated withdrawal and are necessary for the expression of select somatic behaviors^37,38^. However, activity of these ensembles is transient and tightly coupled to specific motor output such as jumping, whereas withdrawal symptoms persist for hours to days and manifest across multiple behavioral domains. Thus, these studies, which have primarily focused on activity within single regions or circuits, have yet to capture a neural signature that reflects the sustained, multidimensional character of withdrawal and the variability in severity across individuals.

One possibility is that such a signature exists at a more distributed level, particularly given the widespread expression of opioid receptors throughout the brain^18^. In humans, large-scale brain recordings have uncovered withdrawal-related alterations in basal ganglia^39–41^, orbitofrontal cortex^39–41^, cingulate cortex^40,42,43^, insular cortex^15,40,44^, striatum^40,45^, and amygdala^15,44^ function. Complimentary behavioral^3,20,46^, cellular^11,19,21^, and circuit^11,15,19–22^ studies in rodents suggest that withdrawal symptoms may emerge from coordinated activity across widely distributed brain networks that integrate neurophysiological changes on both rapid and extended timescales. Currently, it has not been established whether such large-scale neural dynamics exist as a coherent, measurable state, how they evolve over time, or whether they influence the motivational processes driving drug seeking.

Here, we used multiregional electrophysiological recordings and machine learning approaches to identify widespread neural networks associated with fentanyl withdrawal in mice. We first uncovered patterns of neural activity that changed across multiple days of fentanyl use. We then trained a distinct model to predict withdrawal states during precipitated withdrawal and asked whether these learned network patterns generalized to independent animals and contexts. Finally, we examined whether withdrawal-related network activity during voluntary fentanyl self-administration was associated with the escalation of drug intake. Together, these experiments reveal a distributed neural network associated with opioid withdrawal and suggest that the evolution of this network across repeated exposures is linked to individual differences in the escalation of opioid use.

## RESULTS

### Repeated fentanyl exposure alters widespread brain connectivity

To characterize how repeated opioid administration alters widespread neural activity, we implanted male and female C57BL/6J mice with custom multiwire electrode arrays targeting key nodes of corticostriatal, limbic, and midbrain circuits, including the prefrontal cortex (ventromedial orbitofrontal (OFC), infralimbic (IL), prelimbic (PrL), cingulate (Cg)), dorsomedial striatum (DMS), dorsolateral striatum (DLS), nucleus accumbens core (NAcC) and shell (NAcS), bed nucleus of the stria terminalis (BNST), mediodorsal thalamus (mdT), amygdala (AMY), ventral hippocampus (vHip), and ventral tegmental area (VTA). Local field potentials (LFPs) and cellular activity were simultaneously recorded in all sites as mice were exposed to repeated doses of fentanyl across multiple days (**Figure 1A**).

**Figure 1.**
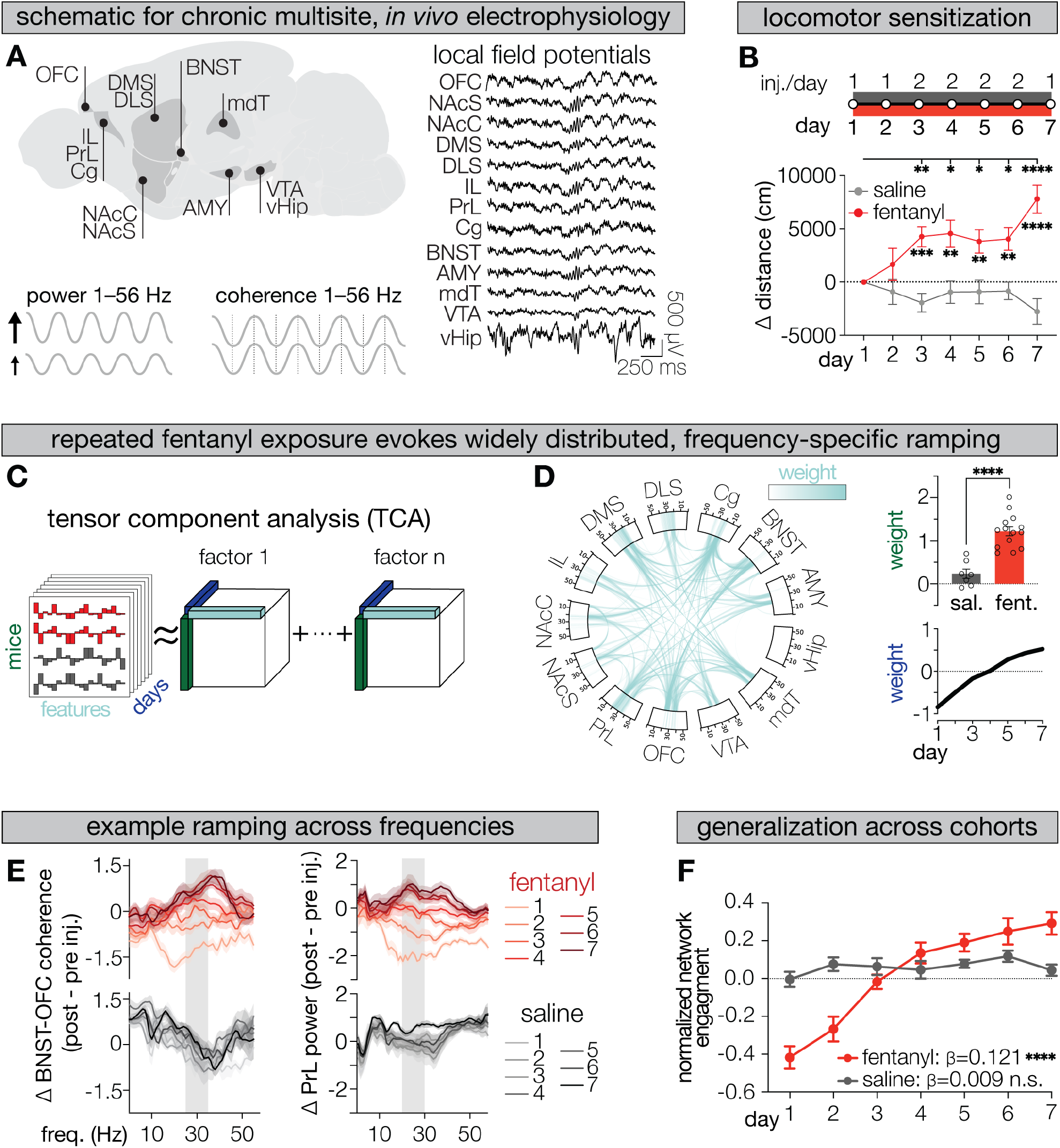
Repeated fentanyl exposure induces progressive, widespread high-beta and low-gamma oscillations. **(A)** Brain regions targeted for *in vivo* local field potential (LFP) recordings (top left) and representative traces (right). LFP spectral features, including within-region power and inter-regional coherence from 1–56 Hz (bottom left) were calculated for each dosing session. **(B)** Experimental design. Mice received once- or twice-daily i.p. injections of fentanyl (5mg/kg; N=14) or saline (N=7) injections for 7 days. Locomotion relative to day 1 showed a significant group effect (*p*<0.0001) and day × group interaction (*p*<0.0001) but no main effect of day (*p*=0.1588; mixed-effects model). Fentanyl-treated mice show significantly greater locomotion compared to saline controls on days 3–7 (*p*<0.05; Bonferroni-corrected, stars below the red line). Fentanyl-induced locomotion on days 3–7 increased significantly relative to day 1 (*p*<0.05; Bonferroni-corrected, stars along the upper bar). **(C)** Tensor component analysis (TCA) schematic. Data were organized into a three-dimensional tensor (mice × features × dosing days) and decomposed into factors capturing shared temporal patterns across mice, features, and days. **(D)** A fentanyl-specific TCA factor characterized by high beta (20–30 Hz) and low gamma activity (30– 40 Hz; left) that ramps across days (right bottom) and has significantly greater expression in fentanyl-versus saline-treated mice (*p*<0.0001; two-tailed, Welch’s *t*-test), as indicated by elevated model-derived weights. Spectral power features are along the rim of the Circos plot, and coherence features connect regions across the plot. Opacity reflects feature weight magnitude. **(E)** Representative baseline-corrected features across fentanyl- and saline-treated mice for BNST-OFC coherence (left) and PrL power (right). Shaded regions highlight prominently represented features captured by the factor in **D**. Ramping across days (lighter to darker traces) can be observed in fentanyl-but not saline-treated mice. **(F)** In a holdout dataset where each mouse’s neural activity was projected into the fixed network from Fig. 1D, network engagement significantly ramped across days in fentanyl- (N=17; *p*<0.0001) but not saline-treated mice (N=12; *p*=0.210; OLS regression). Bar and line graphs depict mean +/-sem. * *p*<0.05, ** *p*<0.01, *** *p*<0.001, **** *p*<0.0001.

Repeated opioid exposure induces progressive neuroadaptations that manifest as behavioral sensitization (i.e. an enhanced locomotor response to the same drug dose over time)^68–70^. To assess this, mice were injected daily with fentanyl (5 mg/kg) or saline for seven consecutive days, and locomotor activity was quantified relative to the first injection day. We observed significant main effects of group (*p*<0.0001) and group × day interaction (*p*<0.0001), but no main effect of day (*p*=0.1588; mixed-effects model; **Figure 1B**). Post hoc analyses revealed that fentanyl-treated mice exhibited significantly greater locomotion than saline controls from days 3–7 (*p*<0.05) and that fentanyl-induced locomotion^68,69^ increased significantly relative to day 1 across all subsequent sessions, with maximal activity observed on day 7 (p<0.05; Bonferroni-corrected). No differences across any days were observed in saline-treated mice.

Next, to determine how repeated fentanyl exposure alters large-scale neural connectivity, we computed spectral power and inter-regional coherence features across all recorded regions and dosing days. To account for individual variability in baseline neural activity, each mouse’s pre-injection responses were subtracted from their post-injection average.

Given that we are primarily interested in identifying patterns of neural activity that changed together across days of fentanyl exposure, we applied an unsupervised dimensionality-reduction approach termed tensor component analysis (TCA; **Figure 1C**). TCA is conceptually similar to principal component analysis (PCA) but extends to higher dimensional data (e.g. mice x LFP features x days). TCA decomposes a data tensor into a set of factors, each defined by: a spatial map of LFP features that co-vary together (a putative network); a temporal profile describing how that network evolves across days (e.g. stable, ramping, or habituating); and a per-mouse loading reflecting the degree of network expression per mouse. Importantly, unlike PCA, where observations such as timepoints or trials are treated as independent samples, TCA explicitly models days as a structured dimension, allowing temporal dynamics to be captured as part of the decomposition rather than collapsed across observations. This approach therefore preserves relationships across mice, features, and days simultaneously, providing an interpretable method to uncover patterns of neural activity that change together over repeated observations and with minimal *a priori* assumptions about the temporal structure of the data.

Our TCA implementation was tuned to uncover the optimal number of components per model, repeated 1,000 times with random initializations, and then clustered to identify highly reproducible (observed in ≥90% of replicates) and stable (≥90% feature similarity) network components (**Figure S2A–B**). This led to a single discovered factor that was uniquely expressed in fentanyl-treated mice, showing markedly higher network expression compared to saline controls (t=8.210, df=15.77, *p*<0.0001; two-tailed Welch’s *t*-test; **Figure 1D**; top right). In fact, saline-treated mice’s average network expression was indistinguishable from zero (*p*=0.375), unlike fentanyl-treated mice (*p*=0.0001; two-tailed Wilcoxon signed-rank test; **Figure 1D** top right), suggesting discovery of a fentanyl-specific network. This network was dominated by high beta (20–30 Hz) and low gamma (30–40 Hz) power and coherence across nearly all recorded regions (**Figure 1D**, left, **S2C–D**). The network’s temporal profile revealed progressive, monotonic increases in expression across days (**Figure 1D**, bottom right). Representative baseline-corrected spectra illustrate this effect, showing clear day-to-day increases in both inter-regional coherence (**Figure 1E** left) and local power (**Figure 1E** right) in fentanyl-but not saline-treated mice.

Large datasets, especially when collected from highly colinear and dependent observations such as multi-site LFP recordings are prone to overfitting, making it challenging to relate physiology to behavior. As such, we reasoned that if the learned connectivity pattern from this rank was truly reflective of the underlying network changes that occur in response to repeated fentanyl, it should be able to classify whether a new set of mice were treated with fentanyl or saline. Therefore, we implanted a new cohort of male and female mice and tested whether this learned connectivity pattern generalized to independent animals. Indeed, a linear mixed-effects model with mouse as a random effect revealed a significant Day × Treatment interaction (*β*=-0.109±0.014, z=-7.70, *p*<0.0001; **Figure 1F**). Post hoc regressions confirmed a significant increase in network activity across days in fentanyl-treated mice (*β*=0.112±0.009, t=9.78, *p*<0.001), but not saline controls (*β*=0.003±0.007, t=0.39, *p*=0.70).

Together, these results reveal a reproducible fentanyl-specific network characterized by progressive increases in high beta and low gamma activity across distributed corticolimbic and striatal regions, reflecting circuit-level adaptations that emerge with repeated opioid exposure.

### A distributed brain network encoding fentanyl withdrawal

Having identified a widespread brain network that emerged through the transition from initial drug use to dependence (**Figure 1**), we next asked how opioid withdrawal affects broad patterns of neural activity.

First, to model withdrawal, fentanyl-dependent mice received naloxone (2 mg/kg), a potent opioid-receptor antagonist that precipitates fentanyl withdrawal (FN; **Figure 2A**). Non-dependent, saline-treated mice continued to receive saline (S) or were given a single injection of fentanyl followed by naloxone (acute fentanyl naloxone, or AFN) to control for handling and the acute pharmacological effects of receptor agonism and antagonism independent of withdrawal (**Figure 2A**). As expected, fentanyl administration increased locomotion, an effect that was exaggerated in dependent animals relative to drug-naïve controls (**Figure 2B**). Naloxone injection sharply reduced locomotion in both FN and AFN groups, often below saline levels (*p*<0.05; Bonferroni-corrected post hoc comparisons within time bins following a significant group x time interaction; **Figure 2B**). Behavioral confirmation of withdrawal was provided by escape jumps, which occurred exclusively in FN mice (FN vs. S *p*=0.0002; FN vs. AFN *p*=0.0033; Dunn-corrected Kruskal–Wallis; **Figure 2C**).

**Figure 2.**
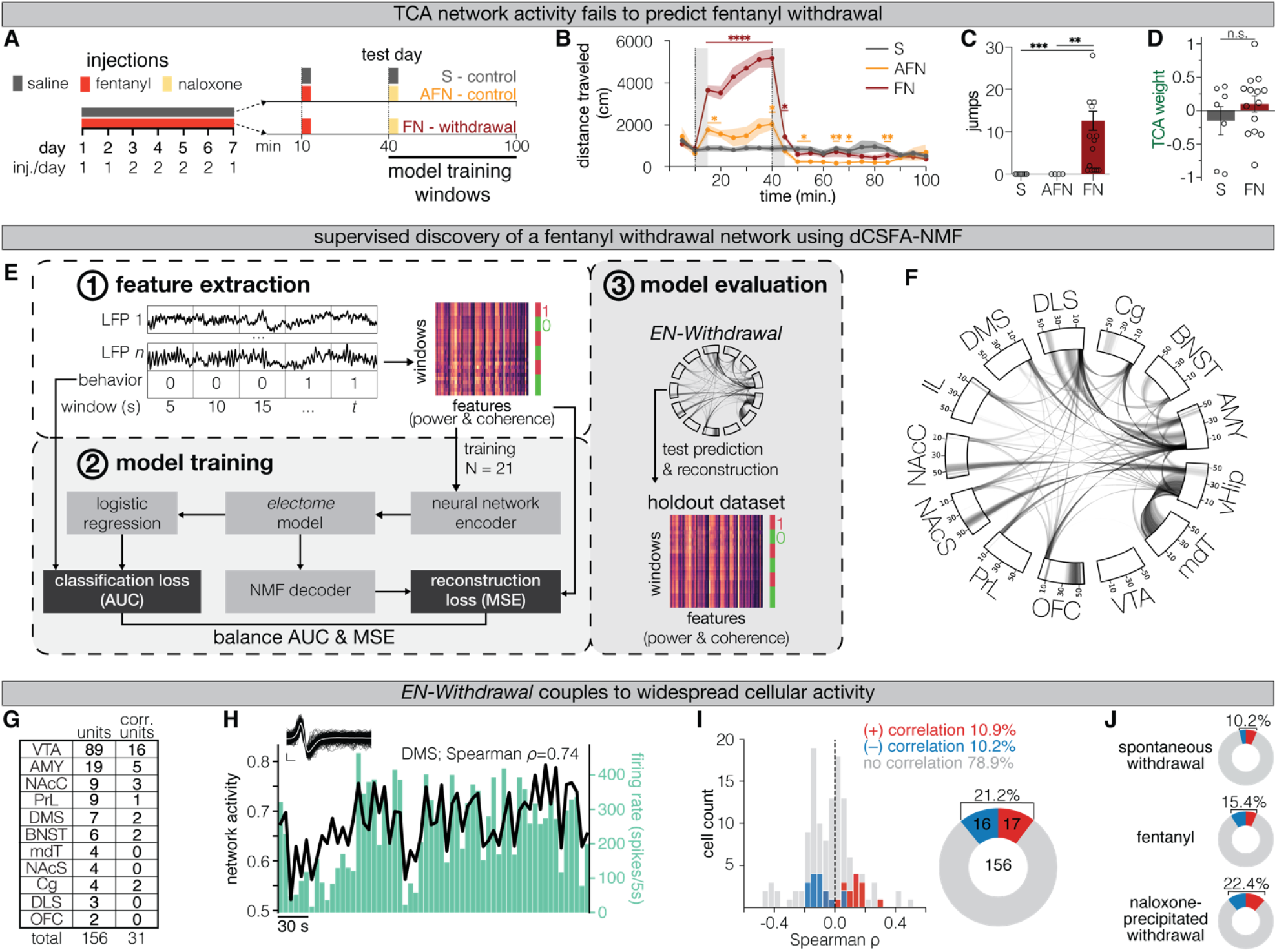
An electome network encoding a fentanyl withdrawal state *(EN-Withdrawal)* reflects distributed cellular activity. **(A)** Experimental design. Following 6 days of fentanyl dosing, mice were tested for withdrawal on day 7. Fentanyldependent mice (FN; N=14) were recorded for 10 minutes to capture spontaneous withdrawal, injected with fentanyl and recorded for 30 minutes, and then injected with naloxone (i.p., 2 mg/kg) to precipitate withdrawal and recorded for an additional 60 minutes. Non-dependent controls received the same dosing schedule (AFN, N=4), or saline only (S, N=7), controlling for pharmacological and handling effects, respectively. **(B)** Locomotor activity on the test day across groups (Time F_2.558, 54.39_=36.06, *p*<0.0001; Group F_2,22_=15.21, *p*<0.0001; Time x Group F_5.116, 54.39_=21.98, *p*<0.0001; mixed-effects model). Bonferroni-corrected comparisons between S vs. FN or S vs. AFN indicated by red or orange stars, respectively. Grey shaded areas denote injection windows. Locomotion is plotted in 5-min bins. **(C)** Only FN mice exhibited withdrawal-induced escape jumps (FN vs. S *p*=0.0002, FN vs. AFN *p*=0.0033; Dunn’s-corrected Kruskal-Wallis). **(D)** Activity of the TCA-based network (Figure 1) in withdrawing (FN) vs. saline control (S) mice (FN vs. S *p*=0.488; Mann-Whitney U). **(E)** Discriminative cross-spectral factor analysis with nonnegative matrix factorization (dCSFA-NMF) workflow. **1**. LFPs were binned into 5-second windows, labeled as withdrawal (1) or non-withdrawal (0), and organized into a two-dimensional array consisting of all time windows across all mice by LFP spectral features. **2**. Features and window labels are used to train a machine learning neural network model that learns an *electome* model, providing class predictions and interpretable neural features used to make those predictions (see Methods for more details). **3**. Predictive and reconstructive accuracy were assessed using held-out datasets not used for training. **(F)** Illustration of the top 15% of neural features in the discovered *electome* network, *EN-Withdrawal*. Spectral power features are along the rim of the Circos plot, and coherence features connect regions across the plot. Opacity reflects feature weight magnitude. **(G)** Table of all isolated and network-correlated units per brain region. **(H)** Representative 5-minute segment of network activity (black trace, left y-axis) and corresponding firing rate of a dorsomedial striatum unit (teal bars, right y-axis; ρ=0.74, *p*<0.0001; two-tailed Spearman). **(I)** Distribution of Spearman correlation coefficients between cellular firing rates and *EN-Withdrawal* activity across the multi-regional population of cells. Units were classified as positively or negatively correlated with network activity using a two-tailed permutation test (*p*<0.05). Distribution of coupled units was significantly greater than expected (*p*<0.0001 twotailed binomial test). **(J)** Proportion of units positively and negatively correlated with network activity during spontaneous withdrawal (*p*=0.0084), fentanyl exposure (*p*<0.0001), and during naloxone precipitated withdrawal (*p*<0.0001; two-tailed binomial test). Bar and line graphs depict mean +/-sem. * *p*<0.05, ** *p*<0.01, *** *p*<0.001, **** *p*<0.0001.

With behavioral confirmation of opioid withdrawal, we first asked whether the same latent activity patterns that distinguished fentanyl- and saline-treated mice during repeated drug exposure also persisted throughout withdrawal. Network activity no longer differed between groups during withdrawal (*p*=0.488; Mann-Whitney U; **Figure 2D**), despite robust separation immediately prior to naloxone administration (day 7, **Figure 1F**). These findings suggest that naloxone rapidly abolished the fentanyl-associated network state identified by TCA.

We next sought to identify the underlying brain network signature of a withdrawal state. To this end, we trained a supervised *binary classifier*, dCSFA-NMF (discriminative cross-spectral factor analysis with nonnegative matrix factorization), to determine withdrawal versus non-withdrawal states for all time windows (**Figure 2E**). While the unsupervised TCA implementation in Figure 1 excels at uncovering how neural trajectories *unfold over time*, identified activity patterns are inherently limited to those that dominate variance. In contrast, dCSFA-NMF is a supervised framework explicitly optimized to uncover static network representations that distinguish between predefined experimental conditions such as withdrawal or non-withdrawal states. This enables optimal discovery of specific state-relevant brain networks, even when such networks explain a small percentage of neural activity. This is particularly relevant because although much of the variance in neural activity can be explained by ongoing behavior^71^, and behavior is often used to infer internal states, the behavior itself is *not* the state^72^. For example, while a sick mouse may exhibit blunted locomotion, this behavior is not sickness in and of itself. dCSFA-NMF has demonstrated strong performance on comparable state classification tasks, such as distinguishing aggressive states *prior* to attack onset^65^ or identifying trait vulnerability to stress in stress-naïve mice^26^. These precedents highlight its ability to uncover latent internal states that persist across time and generalize beyond momentary behavior. This is critical for modeling withdrawal, where outward signs such as jumping or immobility are often highly variable and context specific. Finally, while dCSFA-NMF often predicts with comparable performance of traditional “black box” machine learning models, it retains a high degree of interpretability. This allows us to discover meaningful spectral and regional contributions within the resulting network, providing a biologically grounded and predictive electome model of opioid withdrawal.

Briefly, spectral LFP power and coherence features were segmented into 5-second windows and input into a neural-network-based autoencoder. The encoder learns how to map the LFP features to the network activity scores while the factor loadings (networks) are learned simultaneously to reconstruct the original LFP features in conjunction with the network activity scores to preserve interpretability. These networks can be thought of as the “decoder” in an autoencoder structure. The model jointly optimizes two objective functions: reconstruction loss (mean-squared error – MSE) and classification loss (binary cross-entropy), balancing interpretability and predictive accuracy, respectively. This yields several latent networks, including one supervised network biased toward withdrawal classification and additional unsupervised networks capturing likely distinct physiological variance. Finally, model performance is evaluated on held-out data (see Methods; **Figure 2E**).

The final training dataset included 14 FN, 7 S, and 4 AFN mice, and training cross-validation identified a model with five networks as optimal for maximizing stability, predictiveness, and interpretability (**Figure S3B**). A final model with all training mice and five networks was then fit. Feature weights from the supervised network can be visualized across all power and coherence spectra (**Figure S4**) or can be summarized in a Circos plot that illustrates the top 15% of all features (**Figure S4**, bottom left; **Figure 2F**).

Notably, the strongest feature within this supervised network was orbitofrontal cortex (OFC) gamma-band power (∼40–50 Hz), embedded within a broader network characterized by slow-frequency (delta–theta) inter-regional coherence and faster, region-specific oscillatory power, as observed in the OFC, NAcS, NAcC, DLS, and Cg. When quantifying how often each brain region appeared as a relevant power- or coherence-related feature across frequency bands (delta (1–4 Hz), theta (4–12 Hz), beta (12–30 Hz), and gamma (30–56 Hz)), the DLS and AMY emerge as frequently represented network nodes (**Figure S5A**). While these descriptors help conceptualize the network architecture, it is important to note that the learned network is not a reduced, low-dimensional feature space, but instead leverages *all* input feature weights to define its supervised embedding (**Figure S4**).

This electome network, hereafter referred to as ***EN-Withdrawal***, represents a distributed brain network encoding the neural state of fentanyl withdrawal.

### Widespread cellular activity couples to EN-Withdrawal

We first asked whether the *EN-Withdrawal* network reflected genuine neural population dynamics rather than an abstract statistical construct. To test this, we examined single- and multi-unit spiking activity recorded simultaneously with LFPs across all recorded sites (**Figure 2G**). Unit–network coupling was quantified using a permutation-based approach (**Figure S3C**) that controls the false-positive rate to ≤5% under the null hypothesis of chance alignment. To control for the state of the mice, shuffling occurred only within similar segments (when mice were in spontaneous withdrawal, following fentanyl injections, or following naloxone-precipitation), and correlations were calculated across the entire recording. Across all regions, 21.1% of recorded units exhibited significant coupling with *EN-Withdrawal* activity (*p*<0.05 using two-tailed permutation testing)—a nearly fourfold enrichment over chance (*p*<0.0001 two-tailed binomial test; **Figure 2H–I**). Notably, the highest proportion of network-coupled units occurred during naloxone-precipitated withdrawal (**Figure 2J; Figure S3D**), suggesting that neurons optimally engage with the network during intense withdrawal. These findings confirm that *EN-Withdrawal* dynamics arise from coordinated, large-scale cellular activity distributed across multiple brain regions.

### EN-Withdrawal predicts morphine withdrawal

If *EN-Withdrawal* reflects the underlying biological state of opioid withdrawal, network activity should be able to classify withdrawal in an entire new set of mice exposed to a different opioid drug. To test this, we implanted a new cohort of male and female mice and administered morphine (N=10; 50 mg/kg, i.p.) or saline (N=6) following the same dosing schedule used for fentanyl (**Figure 3A**). Morphine-dependent (MN) mice exhibited withdrawal-induced escape jumps during both spontaneous and naloxone-precipitated withdrawal (Time F_1,28_=0.87, *p*=0.358; Group F_1,28_=5.02, *p*=0.033; Time x Group F_1,28_=0.87, *p*=0.3581; mixed-effects model**; Figure 3B**). Although not every mouse jumped in both the spontaneous and naloxone-precipitated periods, all MN mice jumped at least once in either of the two periods. No saline-treated mice jumped during either period.

**Figure 3.**
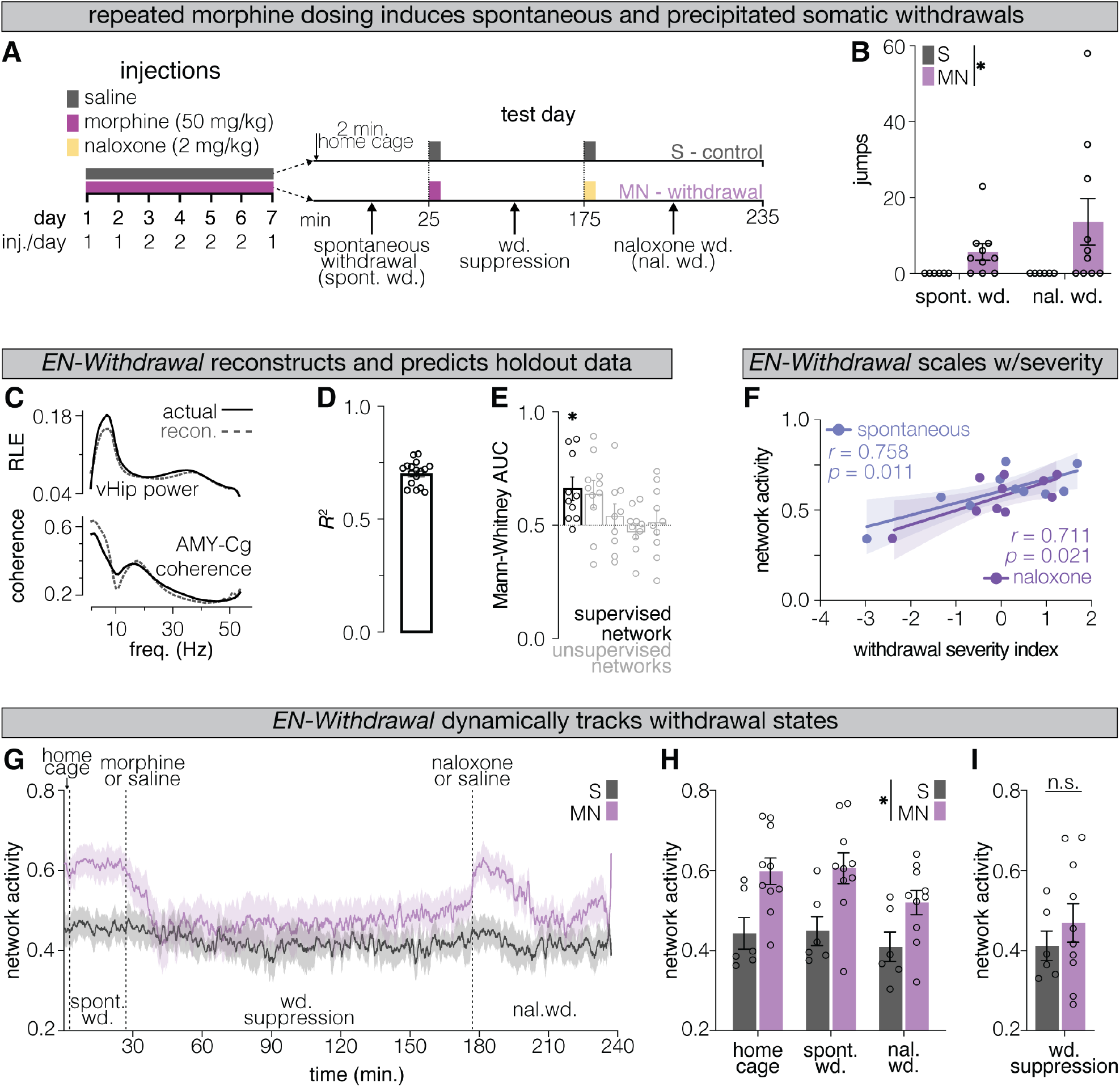
*EN-Withdrawal* activity encodes morphine withdrawal and scales with withdrawal severity. **(A)** Experimental design. Mice received morphine or saline once or twice daily for 6 days. On day 7, morphine-dependent mice (MN) were recorded during spontaneous withdrawal for 25 min, injected with morphine and recorded for 150 minutes, then injected with naloxone to precipitate withdrawal and recorded for an additional 60 minutes. Saline controls (S) received saline injections at matched time points. **(B)** MN mice (N=10) displayed both spontaneous and naloxone-precipitated jumps, whereas S mice (N=6) did not (Time F_1,28_=0.87, *p*=0.358; Group F_1,28_=5.02, *p*=0.033; Time x Group F_1,28_=0.87, *p*=0.3581; mixed-effects model). **(C)** Representative feature reconstruction for ventral hippocampal power (top: relative log energy (RLE)) and amygdala–cingulate coherence (bottom) using the *electome* model. **(D)** Explained variance per mouse in reconstructing LFP spectral features. **(E)** The supervised withdrawal network of the *electome* model, but none of the unsupervised networks, accurately classified withdrawal states in new morphine-withdrawing mice (N=10, *p*=0.019; Bonferroni-corrected two-tailed Wilcoxon signed-rank test). **(F)** Average *EN-Withdrawal* network activity positively correlated with the behavioral withdrawal severity index during spontaneous (*r*=0.758, *p*=0.011) and naloxone-precipitated withdrawal (*r*=0.711, *p*=0.021; two-tailed Pearson correlation). **(G)** *EN-Withdrawal* network activity dynamically tracked withdrawal state transitions. **(H)** Network activity was significantly elevated in morphine withdrawal mice compared to saline controls (Time F_1.68, 23.51_=10.46, *p*=0.001; Group F_1, 14_=7.948, *p*=0.0137; Time x Group F_1.68, 23.51_=1.477, *p*=0.2476; RM two-way ANOVA). **(I)** Following pharmacological reversal of withdrawal in morphine-dependent mice, network activity showed no significant differences between MN mice and saline controls (*p*=0.3585; two-tailed Welch’s *t*-test). Bar graphs depict mean +/-sem. For correlation plots, the line represents the best linear fit with the shaded region indicating the 95% confidence interval. * *p* < 0.05.

The dCSFA-NMF model successfully reconstructed LFP features in this new cohort (**Figure 3C**), with a mean *R-* of ∼70% across mice (**Figure 3D**). The supervised network, but none of the four unsupervised networks, accurately classified withdrawal states in MN mice (AUC=0.622±0.046, *p*=0.019; two-tailed Wilcoxon signe-drank test, Bonferroni-corrected for five network comparisons; **Figure 3E**). Therefore, to avoid unnecessary multiple comparisons moving forward, only the supervised network was used for predictions moving forward.

As withdrawal severity can differ greatly across individuals, even in laboratory-controlled settings, we sought to determine if greater network activity within individuals correlated with greater behavioral severity. Thus, we established a data-driven withdrawal severity index in an independent cohort of male and female mice (N=19; **Figure S6**). Multiple somatic and exploratory behaviors were quantified in spontaneously withdrawing mice and saline controls, including defecation, forepaw tremors, rearing, grooming, immobility, jumping, orofacial behaviors, and booty dragging (**Figure S6B–I)**. PCA dimensionality reduction of z-scored behavioral measures revealed a single behavioral axis (PC1) that explained 54.2% of the variance and clearly separated withdrawing and non-withdrawing mice (*p*<0.0001; Welch’s *t*-test; **Figure S6J–K**). PC1 loadings, which we define as the withdrawal severity index, captured a natural weighting structure in which classical somatic signs (e.g. jumping, orofacial behaviors, tremors, defecation, immobility) contributed positively, while grooming and rearing contributed negatively to the withdrawal severity index (**Figure S6L**).

Strikingly, although *EN-Withdrawal* was trained solely to classify withdrawal states, its activity significantly correlated with behavioral severity during both spontaneous (*r*=0.758, *p*=0.011) and naloxone-precipitated withdrawal (*r*=0.711, *p*=0.021; two-tailed Pearson correlation; **Figure 3F**). While this observation may suggest that *EN-Withdrawal* activity encodes withdrawal severity, it also raises the possibility that activity in the network might simply reflect the specific behaviors expressed in a withdrawal state (e.g., a neural signature of jumping). To evaluate this, we applied the permutation-based statistical approach used to assess unit-network coupling (**Figure 2G–J, S3C–D**) to test for significant behavior–network correlations. For each behavior, the true group-level, two-tailed Spearman correlation between its time course and concurrent network activity was compared to a shuffled null distribution and adjusted with a Bonferroni correction (**Figure S7D–E**). None of the scored behaviors showed significant moment-to-moment coupling with *EN-Withdrawal* activity (**Figure S7F**), indicating that instantaneous changes in network activity are not related to ongoing behaviors.

We next examined how network activity evolves across distinct behavioral phases. Because saline-treated mice have only one state label (non-withdrawal), per-mouse AUCs (**Figure 3E**) reflect within-subject classification of withdrawing mice only, rather than enabling comparisons across groups. To directly compare *EN-Withdrawal* network levels between groups, we projected the neural activity of all mice onto the learned network, yielding a continuous measure of *EN-Withdrawal* activity in MN and S mice across four distinct phases: home cage, spontaneous withdrawal, morphine-induced withdrawal suppression, and naloxone-precipitated withdrawal (**Figure 3A, G**). First, we asked whether network activity was significantly elevated in MN compared to S mice during withdrawal contexts. As expected, *EN-Withdrawal* activity was elevated across home cage, spontaneous, and naloxone-precipitated withdrawal contexts (Time F_1.68, 23.51_=10.46, *p*=0.001; Group F_1, 14_=7.948, *p*=0.0137; Time x Group F_1.68, 23.51_=1.477, *p*=0.2476; RM two-way ANOVA; **Figure 3H**). We then asked whether *EN-Withdrawal* could track a causal pharmacological manipulation of the withdrawal state itself by injecting morphine into withdrawing MN mice. Morphine administration alleviates opioid withdrawal, allowing us to test whether network activity is suppressed accordingly in MN mice. In line with this prior, network activity was statistically indistinguishable from that of saline controls (*p*=0.3585; two-tailed Welch’s *t*-test; **Figure 3I**). Next, we asked whether network activity was elevated compared to that of saline controls 24 hours and 5 days later. Given established time courses of opioid withdrawal in mice, which show heightened withdrawal severity 24 hours following opioid cessation that largely resolves within 5 days^3^, we reasoned that network activity would remain elevated at the earlier, but not the later time point. We found that network activity was greater in MN mice than the saline controls at 24 hours (*p*=0.028; two-tailed Welch’s *t*-test; **Figure S7B**), but not 5 days after the last morphine injection (*p*=0.473; two-tailed Welch’s *t*-test**; Figure S7C**). Together, these causal pharmacological manipulations to suppress (morphine) or induce (naloxone) withdrawal, combined with the persistence of network activity beyond acute manipulations, demonstrate that *EN-Withdrawal* activity is causally and dynamically modulated by withdrawal state.

These results were corroborated in another independent holdout cohort of male and female mice spontaneously withdrawing from morphine following an escalating dosing paradigm (**Figure S8A**; same mice as in Figure S6). After observing that *EN-Withdrawal* could reconstruct this new input data with similar accuracy, ∼70% across animals, (**Figure S8B–C**) we tested our *a priori* hypothesis that network activity would be elevated in withdrawing mice and that activity levels would track individual differences in withdrawal severity. Indeed, significantly elevated activity in withdrawing compared to non-withdrawing mice (*p*=0.0219; one-tailed Welch’s *t*-test; **Figure S8D**). Lastly, across all mice, *EN-Withdrawal* activity positively correlated with behavioral severity (*r*=0.43, *p*=0.032; one-tailed Pearson correlation; **Figure S8E**).

Finally, to test whether *EN-Withdrawal* encodes a withdrawal-specific, rather than a general aversive state, we leveraged the acute noxious response of opioid reversal (**Figure S9A**). Drug-naïve mice (N=8; 4 male, 4 female) developed a conditioned place aversion after a single opioid reversal event via acute fentanyl-naloxone pairing (*p*<0.0001; pre vs. post two-tailed paired *t*-test; *p*=0.0001; one-sample *t*-test; **Figure S9B**), confirming the aversive nature of this manipulation. Therefore, if *EN-Withdrawal* encoded general aversion, network activity should be significantly elevated following opioid reversal compared to a baseline state. However, in an independent holdout cohort of drug-naïve mice (N=6; 3 male, 3 female), acute fentanyl-naloxone pairings had no effect on *EN-Withdrawal* activity relative to baseline or fentanyl-only injections (*p*=0.210; repeated-measures one-way ANOVA; **Figure S9C–D**).

Together, these results demonstrate that *EN-Withdrawal* generalizes across individuals and opioids, predicts withdrawal states and the behavioral severity of withdrawal, and selectively encodes a withdrawal-specific internal state rather than specific behavioral expression or aversion.

### Daily EN-Withdrawal activity predicts urgency and magnitude of fentanyl consumption

Thus far, *EN-Withdrawal* activity has been assessed following *involuntary* opioid exposure and consequently, its relevance to *voluntary* drug taking is unknown. To address this, we implanted a new cohort of male and female mice (N=22) with multisite electrodes and intravenous catheters to chronically record network activity throughout fentanyl intravenous self-administration (IVSA). Mice underwent 22 days of IVSA, beginning with 8 days of continuous-access fixed ratio 1 (FR1) training, followed by 14 days of intermittent-access (IntA) FR1 sessions (**Figure 4A**). During training, each active lever press delivered ∼0.5 µg/kg of fentanyl. During IntA, fentanyl was accessible during a 6-minute window every 30 minutes across 10 consecutive blocks, yielding 1 hour of access distributed across a 5-hour session (**Figure S10A**). Unlike standard paradigms that require lever discrimination and responding criteria to move from training to maintenance phases, we wanted to ensure standardization of duration of access rather than consumption. Thus, all mice completed identical training and IntA schedules with no infusion limits.

**Figure 4.**
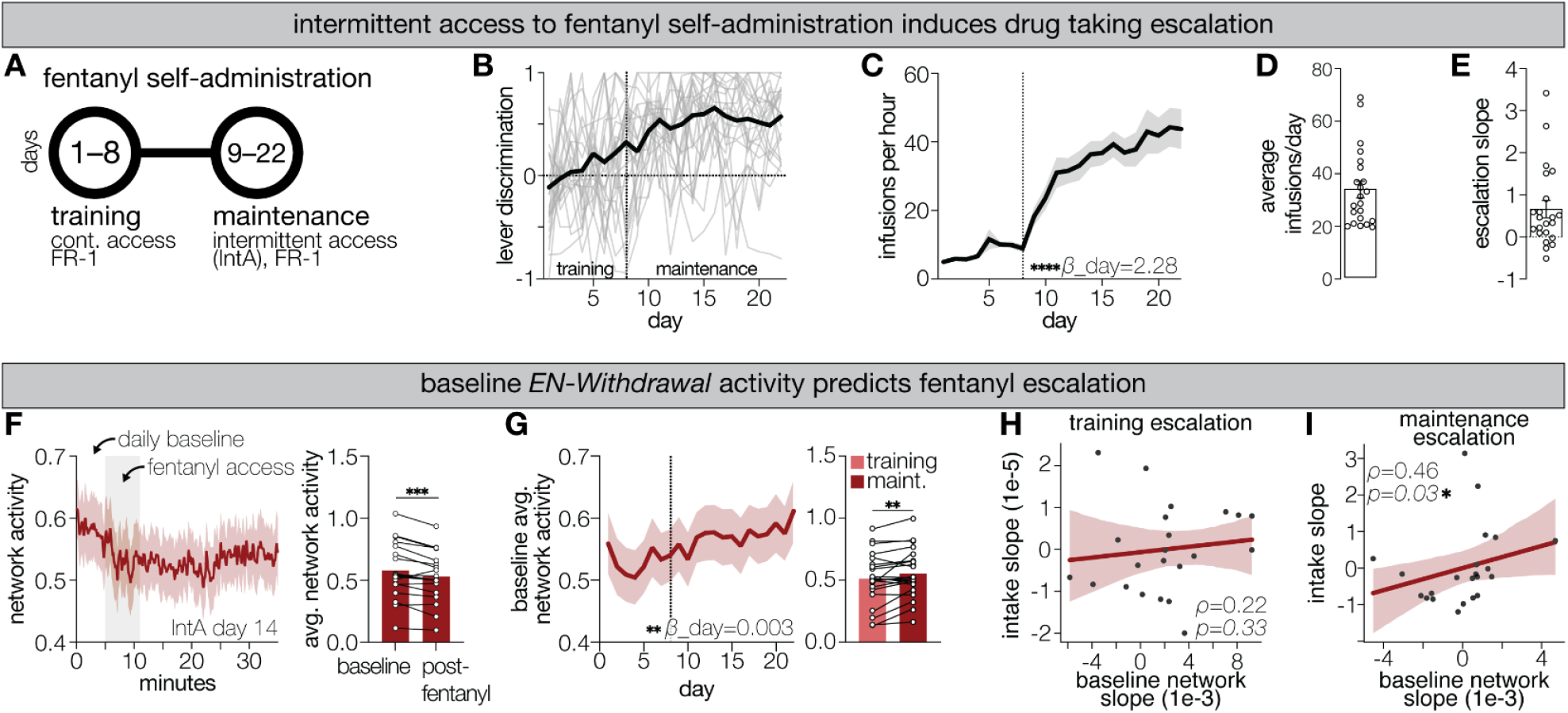
*EN-Withdrawal* predicts daily fentanyl self-administration. (**A**) Timeline for fentanyl self-administration paradigm. (**B**) Lever discrimination index indicating active (1) or inactive (-1) lever preference across days. Light gray lines show individual mouse performance (N=22); solid black line shows group average. (**C**) Number of infusions per hour of fentanyl access across days. Vertical dashed line indicates the last day of training sessions. Mice show significant escalation across days (*p*<0.0001; mixed-effects model). (**D**) Average number of fentanyl infusions per day. (**E**) Escalation slopes per mouse across the full 22-day paradigm. (**F**) Day 14 IntA data projected into *EN-Withdrawal*. Post-fentanyl access period exhibited significantly lower network activity compared to baseline, pre-fentanyl access period (*p*=0.0004; two-tailed paired *t*-test). (**G**) Baseline network activity, taken during the first 5 minutes of each session, escalated across days (*p*=0.0012; mixed-effects model) and was significantly elevated during the maintenance phase compared to the training phase (*p*=0.0091; two-tailed paired *t*-test). (**H**) Escalation in fentanyl intake did not correlate with escalation in network scores during training (ρ=0.22, *p*=0.33; two-tailed Spearman correlation). (**I**) Escalation in fentanyl intake significantly correlates with escalation in network scores throughout the maintenance phase (ρ=0.46, *p*=0.03; two-tailed Spearman correlation). Bar graphs depict mean +/-sem. For correlation plots, the line represents the best linear fit with the shaded region indicating the 95% confidence interval. * *p*<0.05, *\*\* p*<0.01, *\*\*\* p*<0.001, *\*\*\*\* p*<0.0001.

Lever discrimination during training was initially poor but improved rapidly during IntA, with mice showing a clear preference for the active lever (Day F_21,462_=18.02, *p*<0.0001; Lever F_1,22_=43.28, *p*<0.0001; Day x Lever F_21,439_=11.29, *p*<0.0001; mixed-effects model; **Figure 4B**, significant Bonferroni-adjusted post hoc comparisons indicated in **Figure S10B**). Consistent with prior reports showing that IntA promotes escalation of opioid intake^7^, fentanyl consumption increased progressively across days (β_Day=2.28 ± 0.32, z=7.09, *p*<0.0001; linear mixed-effects model with a random intercept and slope for individual mice; **Figure 4C**). Unsurprisingly, substantial variability emerged across individuals’ daily infusion counts (**Figure 4D**) and escalation rates (**Figure 4E)**. Nonetheless, we observed a significant reduction in *EN-Withdrawal* activity during the post-fentanyl access period compared to the baseline pre-access period on the last day of self-administration testing (*p*=0.0004; two-tailed paired t-test; **Figure 4F**). Additionally, baseline network activity progressively increased across days (β_Day=0.003 ± 0.001, z=2.88, *p*<0.004; linear mixed-effects model with a random intercept and slope for individual mice; **Figure 4G**) and was significantly elevated during the latter maintenance phase compared to early training days (p=0.0091; two-tailed paired t-test; **Figure 4H**), indicating an accumulation of network activity independent of immediate drug availability.

Given pronounced individual variability in both fentanyl and network escalation rates, we asked whether the rate of increase in *EN-Withdrawal* activity, measured at the start of each session before any drug exposure for the day, covaried with the rate of fentanyl intake escalation, and whether this relationship depended on prior drug experience. For each mouse, we extracted empirical Bayes estimates of individual escalation rates (random slopes) from a joint mixed-effects model fit across all sessions. During the training phase, when animals are not experienced users, no significant relationship was observed between network and fentanyl escalation rates (ρ=0.22, *p*=0.33; two-tailed Spearman correlation; **Figure 4H**). In contrast, during the maintenance phase, *EN-Withdrawal* escalation rates significantly predicted fentanyl escalation (ρ=0.46, *p*=0.03; two-tailed Spearman correlation; **Figure 4I**), indicating that mice with steeper increases in baseline *EN-Withdrawal* activity also escalated fentanyl consumption more rapidly. These findings demonstrate that the relationship between withdrawal network dynamics and voluntary fentanyl escalation is experience-dependent, emerging selectively after prolonged opioid exposure when negative reinforcement has likely become a dominant motivational force.

## DISCUSSION

In this study, we set out to discover how opioid withdrawal is encoded by the brain, and whether such encoding shapes drug seeking. Using multisite LFP recordings across 13 brain regions combined with interpretable machine learning, we identify two distinct network signatures of opioid experience. The first, uncovered via TCA, is a progressive increase in inter-regional high beta and low gamma coherence that ramps monotonically across days of repeated fentanyl exposure, paralleling the development of locomotor sensitization. The second, and central contribution of this work is a distributed network characterized by regional gamma oscillations and slow interregional coherence that encodes the internal state of opioid withdrawal. Activity in this network encodes the behavioral severity of withdrawal and increases in the activity of this network across days predicts fentanyl intake escalation in experienced animals. Together, these findings establish a systems-level neurophysiological substrate for the negative reinforcement cycle of addiction.

### Repeated opioid exposure drives large-scale connectivity changes

Withdrawal emerges only after opioid exposure that is sufficient to produce dependence, yet how repeated use reshapes large-scale communication across the brain remains poorly understood. Using unsupervised TCA, we identified a monotonic increase in high beta and low gamma coherence across fentanyl exposure (**Figure 1**), which generalized across individuals, reinforcing the link between repeated opioid exposure and distributed neural adaptations.

Although a single, ramping connectivity pattern was consistently and uniquely observed in fentanyl-treated mice across both training and test sets, TCA is optimized to isolate stable, recurrent covarying activity patterns rather than to exhaustively capture all ongoing neural dynamics. Therefore, it is conceivable that this factor likely represents only a subset of the underlying adaptations. Nonetheless, the consistent emergence of this ramping pattern across model training repeats and cohorts highlights a robust and generalizable signature of network-level plasticity induced by repeated opioid exposure, supporting the view that dependence arises from widespread circuit-level remodeling beyond localized receptor adaptations.

### A distributed electome encodes withdrawal states

While prior studies have provided critical insights into regional neural changes during withdrawal, our findings extend this work by revealing how these localized dynamics integrate within a distributed, multiscale network that both encodes and predicts withdrawal states. Remarkably, *EN-Withdrawal* generalized to independent cohorts, accurately predicting both spontaneous and naloxone-precipitated withdrawal in morphine-dependent mice, even under distinct dosing regimens (**Figure 3, Figure S8**). This generalizability suggests that the withdrawal state, regardless of opioid type or dosing pattern, has a conserved neurophysiological signature. By capturing fast neural oscillations in the context of slow behavioral timescales (withdrawal developing over days), we bridge temporal scales to identify how transient neural events accumulate onto a convergent network state.

These findings yield several noteworthy insights linking withdrawal to internal brain states. First, *EN-Withdrawal* activity was elevated regardless of the variation in ongoing, overt somatic signs (**Figure S7D–F**). This demonstrates that the network captures a state that is not simply reactive to motor activity. Moreover, while the behavioral expression of spontaneous and naloxone-precipitated withdrawal are known to differ^22^, *EN-Withdrawal* activity correlated with overall behavioral severity across both forms, decoupling network activation from any single behavior or dosing regimen. Second, *EN-Withdrawal* activity remained elevated when animals were transitioned from their home cage to the behavioral monitoring arena, demonstrating that this state representation is stable across contexts and not dependent on external environmental cues. Third, morphine robustly suppressed *EN-Withdrawal* activity for over 150 minutes, while naloxone produced a rapid and sustained increase in activity, consistent with the pharmacological dynamics of withdrawal. Strikingly, *EN-Withdrawa*l activity remained significantly elevated 24 hours after the test session, well beyond the pharmacological window of morphine, but normalized by five days of withdrawal. This timescale of persistence, spanning hours to days, aligns with the temporal profile of withdrawal symptoms in both rodents and humans^3^, and stands in contrast to previously reported withdrawal-related neural signals that extinguish within seconds to minutes^19,41^. Lastly, network activity patterns showed no significant changes following acute fentanyl-naloxone reversal in drug-naïve mice despite this manipulation being sufficient to produce a robust conditioned place aversion with only a single contextual pairing, confirming that the network is unlikely to simply reflect a generally aversive state.

Together, these findings support a novel conceptual framework that directly aligns *EN-Withdrawal* activity patterns with known features of internal states, including persisting across time, scaling with intensity, generalizing across context (and subjects), and reflecting pleiotropic effects^72^. This is, to the best of our knowledge, the first account of a physiological signature of an internal brain state of withdrawal.

### Regional contributions to withdrawal encoding

Within the framework of allostatic models of addiction, in which motivated drug use progresses from reward seeking to withdrawal avoidance to compulsive behavior^12^, several activity patterns identified by *EN-Withdrawal* may help bridge withdrawal states to future compulsive drug use. For example, repeated stimulus–response associations drive habit formation through prefrontal cortical inputs to the DLS^73–77^, a pathway well positioned to link withdrawal-driven behaviors to habitual drug seeking. *EN-Withdrawal* not only identified the DLS as a central hub (**Figure S5A**) and linked its coherent activity with many other regions (**Figure 2F**), but training dCSFA-NMF models on only the DLS and a single other region further identified connections between the anterior cingulate cortex and DLS are predictive circuit components (**Figure S5B–C**).

The amygdala has long been implicated in withdrawal states^21^ and emerged as another densely connected hub within *EN-Withdrawal* (**Figure S5A**). While most prior studies have focused on the central amygdala, our results point towards a critical contribution of the basolateral amygdala in shaping the negative affective components of withdrawal. The basolateral amygdala provides dense inputs to the ventral hippocampus and this circuit has been reported to engage aversive responses and contribute to withdrawal-induced anxiety states^78^. However, given the volume-conducted nature of LFP signals, we cannot rule out the possibility that activity attributed to the BLA may partially reflect engagement of neighboring amygdalar subregions and therefore simply refer to this activity as amygdala activity throughout the study.

Lastly, orbitofrontal cortical (OFC) inputs to the striatum form a core compulsivity circuit implicated in both obsessive–compulsive disorder and excessive drug use in addiction, in both preclinical and clinical populations^79^. Notably, the strongest within-region feature of *EN-Withdrawal* was an increase in OFC gamma power (**Figure 2F**), which may signal ongoing plasticity that predisposes future maladaptive OFC–striatal connectivity. Interestingly, unlike other prefrontal cortical regions, including the prelimbic, infralimbic, and cingulate cortices that showed ramping interregional power activity from repeated fentanyl injections (**Figure 1D–E**), the OFC exhibited little to no increases in power during fentanyl exposure and instead was evoked by withdrawal (**Figure 2F**). This elevated gamma activity could arise from enhanced excitability and firing of parvalbumin-expressing interneurons during withdrawal^4^, which are key regulators of cortical oscillatory synchrony. Such changes could ultimately bias behavior toward compulsive drug use. Future studies should leverage similar forms of chronic *in vivo* monitoring techniques to directly test how withdrawal-induced OFC plasticity contributes to the transition from dependence to compulsivity.

### EN-Withdrawal reflects withdrawal severity

Accurately quantifying withdrawal severity in rodents remains a long-standing challenge, as scoring methods are often non-standardized and rely on arbitrary weighting of behavior^80^. Most studies quantify multiple behavioral phenotypes (e.g. jumping, tremors, or rearing) into a “composite score” by assigning arbitrary, experimenter-defined weights to each behavior before summing them all together^46,58–62^. This approach can allow one behavior to dominate or obscure meaningful group-level variation^80,81^. To address these limitations, we adopted a data-driven approach by quantifying a broad ethogram of established withdrawal-related behaviors and deriving behavioral weights directly from the data using PCA (**Figure S6J**). This analysis revealed a single latent dimension that best separated withdrawing from non-withdrawing mice (**Figure S6K**) and provided empirically defined weights for each behavior (**Figure S6L**). The resulting withdrawal severity index not only circumvents arbitrary weighting but also generalized across independent cohorts, offering a reproducible and quantitative measure of withdrawal severity.

Impressively, beyond simple classification, the magnitude of *EN-Withdrawal* activity scaled with individual behavioral withdrawal severity (**Figure 3F, Figure S8E**). Notably, network activation persisted over extended withdrawal periods despite evolving behavioral manifestations (e.g., early escape jumping versus later inactivity; **Figure S7D–F**), suggesting that *EN-Withdrawal* reflects a sustained internal state rather than overt motor output. Importantly, by deriving a widespread network signature from neural activity rather than relying solely on behavioral scoring, this work advances the field towards capturing an internal state of addiction based on measurable brain physiology^72^.

In a field where diagnostics and classification rarely incorporate direct measures of neural state, our findings provide a precedent for how large-scale brain network dynamics can serve as objective indicators of internal states, augmenting traditional behavioral approaches.

### EN-Withdrawal magnitude predicts escalation of fentanyl use

Voluntary intravenous self-administration (IVSA) has served as the gold-standard rodent model of addiction since the 1960s^82^. Consistent with this literature, we find that mice reliably self-administer fentanyl across multiple weeks and display hallmark addiction-like behaviors, including robust escalation of drug intake (**Figure 4C**). Continuous multisite recordings throughout IVSA provided a unique opportunity to assess whether *EN-Withdrawal* activity was relevant to voluntary drug taking. Although fentanyl use acutely suppressed network activity, in line with withdrawal relief as a reinforcing effect, network activity rebounded and climbed progressively over the course of the experiment, consistent with a dynamically shifting set point, or allostatic state, hypothesized across predominant theories of addiction^12^.

Crucially, it was not absolute network activity, but the rate of network escalation that predicted individual differences in fentanyl intake escalation. This relationship was specific to the drug-experienced maintenance phase and was absent during early training, when animals are likely motivated primarily by fentanyl’s rewarding effects rather than withdrawal avoidance. This experience-dependent emergence of the network-behavior relationship is important as it suggests that *EN-Withdrawal* becomes a meaningful marker of intake escalation only after dependence has developed, consistent with the transition from positive to negative reinforcement that characterizes the progression to addiction. Moreover, the fact that this relationship was captured at the level of individual escalation rates, rather than group averages, underscores its potential relevance as a within-subject biomarker of addiction vulnerability. Together these data link a sustained neural state of withdrawal (*EN-Withdrawal*) to the worsening motivational dynamics that characterize addiction.

### Limitations and future directions

Several limitations of the present study warrant consideration. First, while *EN-Withdrawal* provides a robust framework for understanding the neural architecture of opioid withdrawal, this study does not yet address how network adaptations that arise from repeated opioid exposure (**Figure 1**), rather than during withdrawal, relate to subsequent withdrawal severity. Determining whether individuals showing stronger expression of the fentanyl-specific network also exhibit heightened *EN-Withdrawal* activity and more severe behavioral withdrawal would provide critical insight into how progressive neural adaptations evolve into dependence. Second, while unit–network correlations (**Figure 2**) suggest broad cellular engagement, unit counts are relatively modest. Although traditional neuroscience studies typically utilize direct cellular manipulations via optogenetics or chemogenetics to overcome such shortcomings and provide causal links, the field lacks reliable tools to directly modulate large scale network dynamics like those reported here. Nonetheless, future studies combining targeted perturbations of circuits within the larger network architecture could determine whether modulation of specific cellular ensembles can mediate changes in oscillatory dynamics, network activity, and withdrawal expression. Third, although both male and female mice were included throughout, we did not explicitly test for sex differences; instead, we prioritized discovery of network dynamics that generalized across sex. However, given the well-established influence of sex on addiction-related behaviors^83–85^, future studies should directly investigate sex-specific contributions and potential variability in network expression. Lastly, while we primarily investigated acute withdrawal states marked by somatic symptoms, these findings could benefit from exploration of how *EN-Withdrawal* dynamics evolve during protracted withdrawal phases, in which somatic signs cease but affective, cognitive, and memory-related dysfunction arises^3^.

### Conclusions

This work integrates chronic multisite LFP recordings with interpretable machine learning to identify a novel, widely distributed brain network, *EN-Withdrawal*, that encodes the internal state of opioid withdrawal. *EN-Withdrawal* activity serves as a predictive neuro-biomarker of withdrawal severity that generalizes across sex, opioids, and dosing paradigms. Most importantly, its evolution across time predicts individual fentanyl intake escalation in drug-experienced animals. These findings elucidate a neurobiological substrate for an internal state underlying the negative reinforcement cycle of addiction: a distributed network that becomes active during withdrawal and potentially motivates behaviors aimed at relief. Disrupting this network, or the circuit mechanisms that drive its escalation, may offer a principled target for interventions aimed at breaking the compulsive drug use cycle driven by withdrawal avoidance.

## MATERIALS AND METHODS

***Animal care & use:*** Male and female C57BL/6J (C57) mice (N=107) were purchased from Jackson Labs at 7–10 weeks of age. Unless otherwise specified, mice were housed 3–5 per cage, kept on a 12-hour light/dark cycle, and maintained in a humidity- and temperature-controlled room with water and food available *ad libitum*. Male and female mice, 10–18 weeks old at the start of experimentation, were used for all experiments presented in this study. Across studies, male and female mice were handled by both male and female experimenters. All experimental tests were conducted with approved protocols from the Duke University Institutional Animal Care and Use Committee.

### Electrode implantation surgery

The electrode implantation surgery procedure has been described previously^47,48^. Mice were anesthetized with 1.5% isoflurane, placed in a stereotaxic device and metal ground screws were secured above anterior cranium (left of midline) and cerebellum (midline). A third screw was secured laterally, roughly half-way between the other screws. Thirty-two tungsten microwires were arranged in bundles such that each brain region was targeted with two wires. The electrode was designed to target the basolateral amygdala (AMY), medial dorsal nucleus of thalamus (mdT), nucleus accumbens core (NAcC) and shell (NAcS), ventral tegmental area (VTA), prelimbic cortex (PrL), cingulate cortex (Cg), infralimbic cortex (IL), ventral hippocampus (vHip), ventromedial orbitofrontal cortex (OFC), bed nucleus of the striata terminalis (BNST), dorsomedial striatum (DMS), and the dorsolateral striatum (DLS). Brain targets were centered based on stereotaxic coordinates referenced to bregma (AMY: -1.4 mm AP, 3.0 mm ML, -3.95 mm DV from dura; mdT: -1.58 mm AP, 0.4 mm ML, -3.0 mm DV from dura; VTA: -3.0 mm AP, 0.25 mm ML, -4.25 mm DV from dura; vHip: -3.0 mm AP, 3.0 mm ML, -3.75 mm DV from dura; PrL: 1.62 mm AP, 0.25mm ML, 1.65 mm DV from dura; Cg: 1.62 mm AP, 0.25mm ML, 1.05 mm DV from dura; IL: 1.62 mm AP, 0.25mm ML, 2.25 mm DV from dura; NAcC: 1.1 mm AP, 2.25 mm ML, -3.5 mm DV from dura, implanted at an angle of 16°; NAcS: 1.1 mm AP, 2.25 mm ML, -4.0 mm DV from dura, implanted at an angle of 16°; OFC: 2.5 mm AP, 1.25 mm ML, -2.0 mm DV from dura; BNST: 0.1 mm AP; 0.8 mm ML; 4.5 mm DV from dura; DMS: 1.5 mm AP, 2.25 mm ML, 1.7 mm DV from dura, implanted at an angle of 16°; DLS: 1.5 mm AP, 3.25 mm ML, 1.7 mm DV from dura, implanted at an angle of 16°).

All mice were additionally targeted with two wires in the central amygdala and anterior insula each, but analyses of these regions were left out of this work due to inconsistent targeting across mice. The periaqueductal gray was targeted in about three-quarters of the mice here but was also excluded from analysis. All bundles were implanted in the left hemisphere. Each electrode bundle was coated with a fluorescent dye (NeuroTrace™ CM-DiI Tissue-Labeling Paste; ThermoFisher #N22883) immediately prior to implantation to aid in postmortem identification of electrode tracks. A metal ground wire was secured to the anterior and posterior screws, and the implanted electrodes were anchored to all three screws using dental acrylic. To mitigate pain and inflammation related to the procedure, all animals received carprofen (5 mg/kg s.c.) once prior to surgery and every 24 hours following for 72 hours.

### Jugular catheterization

Between 7–10 weeks old, adult male and female C57BL/6J mice (N=22) were implanted with indwelling catheters (Instech Laboratories, Inc., Plymouth Meeting, PA) into the right jugular vein^49^. First, polyurethane catheters with a bead 1.2 cm from the catheter tip (part #VAB62BS/25; Instech Laboratories, Plymouth Meeting, PA) were prepared by cutting excess tubing ∼3 cm from the bead and affixed to a vascular access button (part #VAB62BS/25; Instech Laboratories, Plymouth Meeting, PA). Then, mice were anesthetized with 1.5% isoflurane and placed on their backs with their noses fixed into a nose cone. Incisions were made above the right pectoral muscle and mid-scapular line. Then, catheters were threaded from the mid-scapular incision under the skin of the shoulder and out from the pectoral incision. The jugular vein was carefully located, raised up, cut, and the catheter was carefully guided into the vein until the catheter bead reached the jugular incision. The catheter was fixed into place with two sutures around the jugular vein. For 3 days post-surgery, animals received amikacin (10 mg/kg s.c.) to prevent perioperative infection. To maintain patency and prevent infection afterwards, catheters were flushed 1–2 x daily with 50 µL of a 30 U/mL heparin and 1 mg/mL gentamicin solution.

Catheter patency is traditionally tested by infusing small amounts of ketamine into the catheter and testing for rapid sedation^49–51^. However, to avoid known confounds of ketamine effects on neural dynamics^52,53^, all mice were assumed to have patent catheters unless suspicion was raised. Suspicious circumstances include failure to reliably self-administer fentanyl after 4 days of operant training or rapid, sudden reduction in drug-taking behaviors during daily self-administration. No mice used in this study were subjected to ketamine for patency testing.

### Histological confirmation

Histological analysis of implantation sites was performed at the conclusion of all experiments to confirm electrode placement (**Figure S1**). Animals were perfused with 4% paraformaldehyde (PFA) and brains were harvested and stored in PFA for 24 hours. Then, brains were rinsed in PBS 2–4 times, sliced at 40 µm (Leica Vibrating Blade Microtome) and stored in PBS with 0.05 mM sodium azide for longterm storage. Targeted brain slices were chosen and placed into 24-well plates, incubated with nuclear Hoechst (Invitrogen™ DAPI and Hoechst Nucleic Acid Stains; Fisher Scientific #H1399) stain, rinsed 3–5 times with PBS, and then mounted and cover slipped on standard microscope slides for imaging. Slides were imaged at 4x, and occasionally 10x or 20x when higher resolution was needed for electrode placement confirmation, with an Olympus VS200 Slide Scanner. Animals were excluded from analysis if any targeted region was mistargeted or could not be confirmed histologically. In total, 14 mice were excluded, yielding an overall targeting accuracy of ∼88%.

### Fentanyl and morphine dosing protocols

To induce withdrawal, mice underwent one of three opioid dosing protocols indicated throughout the text.

#### 1) Fentanyl Withdrawal

At least two weeks following electrode implantation, mice (N=21) were pseudorandomly assigned to receive either saline (n=7; 4 males, 3 females) or fentanyl (n=14; 8 males, 6 females) injections. Fentanyl (Hikma, Eatontown, NJ 07724, USA, NDC: #0641-6030-01) was purchased prediluted in saline in a 0.5 mg/mL solution and injected intraperitoneally at 10 mL/kg or an equivalent volume of saline. On the first 2 days, mice received a single morning injection while in a clear Plexiglas monitoring box; on the subsequent 4 days, mice received twice-daily injections (morning in monitoring box, evening in home cage). On the 7^th^ day, mice received either saline injections or a combination of fentanyl and naloxone hydrochloride (Akorn Inc. Lake Forest, IL 60045 #17478-0042-10; 2 mg/kg) to induce withdrawal, described in more detail below.

Mice were placed into a dimly lit behavioral room (50 lux) and allowed to habituate for 1 hour prior to behavior recordings. Then, all cage mates were moved into a holding cage and each mouse, one at a time, was connected to the recording system via a head-stage and placed back in its original home cage for 5 minutes to acclimate. LFP and video recordings were collected during morning injections on all experimental days. On day 1, mice were recorded for 2 minutes in their home cage before being moved into a monitoring box for 20 minutes, injected with saline or fentanyl based on their group assignment and then recorded for an additional 20 minutes. On days 2–6 (mornings), mice were recorded for 2 minutes in their home cage, then 5 minutes in the monitoring box, injected with saline or fentanyl based on their group assignments and then recorded for an additional 20 minutes. Evening injections on days 3–6 were done in the home cage without electrophysiology or video collection. On day 7 (test day), following habituation, mice underwent the following:

1. Mice were recorded for 2 minutes in their home cage before being placed in the monitoring box for 10 minutes.
2. Mice were injected with saline or fentanyl based on group assignment and recorded for 30 minutes.
3. Mice were then given a final injection of saline or naloxone based on group assignment and recorded for 60 minutes.
4. A subset of saline mice were rerun the subsequent day and given fentanyl and then naloxone as an acute fentanyl naloxone control (AFN: n=4; 2 males, 2 females) in non-dependent mice (**Figure 2A**).

#### 2) Morphine Withdrawal (Constant)

At least two weeks following electrode implantation, mice (N=16) were pseudorandomly assigned to receive either saline (n=6: 3 males, 3 females) or morphine injections (50 mg/kg; n=10: 3 males, 7 females). Morphine (Hospira, Lake Forest, IL 60045, USA, NDC: # 0641-6127-01 or Hikma, Eatontown, NJ 07724, USA, NDC: #0641-6127-25) was purchased prediluted in saline at 10 mg/mL, further diluted to 5 mg/mL, and injected intraperitoneally at 10 mL/kg. Morphine dosing followed the same timeline as fentanyl dosing: once daily for the first 2 days, followed by 4 days of twice-daily injections and a final injection on the 7^th^ day. Injections on days 1-6 were in the home cage. Electrophysiology and video recordings were only conducted on the test day (day 7). Control mice received only saline injections, matched for timing and volume.

For electrophysiology recordings, mice followed the same habituation protocol described above. On day 7:

A. Mice were recorded for 2 minutes in the home cage, followed by 25 minutes in the monitoring box to allow for observation of spontaneous withdrawal.
B. Mice were injected with saline or morphine, followed by recording for 150 minutes.
C. Finally, mice were injected with saline or naloxone based on group assignment and recorded for an additional 60 minutes (**Figure 3A**).

#### 3) Morphine Withdrawal (Escalating)

Mice (N=23) received twice daily subcutaneous saline (n=8: 4 males, 4 females) or morphine (n=15: 8 males, 7 females) injections at a volume of 10 mL/kg that escalated across 6 days of injections in their home cage as follows: 20, 40, 60, 80, 100, 100 mg/kg^30,54–56^. 12–16 hours following the final morphine injection, mice were recorded for 2 minutes in their home cage followed by 30 minutes in the monitoring box to observe spontaneous withdrawal.

### Conditioned place aversion (CPA)

To test for the acute aversive effects of opioid reversal, mice (N=8: 4 males, 4 females) were tested on a single-pairing, two-chamber conditioned place aversion (CPA) paradigm, adapted from Tian et al., 2022^57^.

A. On the first day, mice were placed in the middle of the two-chamber arena and allowed to freely explore both chambers for 15 minutes. An initial chamber preference was determined by quantifying the amount of time spent in each chamber.
B. On the morning of the second day, mice received an i.p. injection of saline. They were then placed back in their home cage and injected again with saline immediately before being placed into their less preferred chamber for 15 minutes.
C. In the afternoon, mice received an i.p. injection of fentanyl (5 mg/kg), were placed back in their home cage, and then injected with naloxone (2 mg/kg) immediately before being placed into their more preferred chamber for 15 minutes.
D. On the third day, mice were allowed to freely explore both chambers again for 15 minutes.

All sessions were video-recorded in real time using a NeuroMotive (Blackrock Microsystems, Inc., Salt Lake City, UT) camera system for a top-down view of the arena. Mice were tracked using Ethovision X12 (Noldus, Wageningen, the Netherlands) and center point location was used to determine the percent of time spent in each chamber. A CPA is determined if the time spent in the acute fentanyl-naloxone chamber was significantly less than the initial preference for that chamber (post-vs. pre-preference) and if the post-conditioning time spent was significantly below 50% (post-conditioning aversion).

### Fentanyl self-administration

Briefly, mice (N=22) were trained to operantly respond for intravenous delivery of fentanyl for 8 days within a 10-day period, followed by 14 days of intermittent access sessions. Fentanyl was diluted in sterile saline to form a final concentration that would provide a cohort average dose of 0.5 µg/kg/infusion. Self-administration boxes were custom designed in-house but utilized commercial lever apparatuses and syringe infusion pumps (Med Associates, Inc., Fairfax, VT). Active lever assignment was pseudorandomized across mice and indicated by the presence of a cue light directly above the lever which remained on throughout the session; the inactive lever’s light was off throughout the session. Each infusion during training was followed by a 20-second timeout period during which the lever was retracted, such that no additional fentanyl could be administered. Throughout this timeout period, the cue-light flashed at 1 Hz.

No autoshaping sessions (free infusions independent of lever presses) were conducted. All training sessions followed a continuous-access, fixed-ratio 1 (FR1) schedule, meaning that mice had access to fentanyl for the entirety of the session (minus post-infusion timeouts) and each active lever press resulted in an infusion of 18 µL of fentanyl. Mice began training with an overnight, 12-hour extended session with standard laboratory chow provided ad libitum. Mice were removed from the self-administration boxes the following morning and given the day off. The subsequent 3 days consisted of 2-hour training sessions. This 5-day block was then repeated: overnight, off, training day 6, 7, and 8.

While most studies typically aim for specific lever discrimination criteria and minimum and/or maximum infusion rates that must be met to graduate from training, we hoped to leverage the variability in drug seeking and taking strategies across mice. Therefore, all mice moved from training to maintenance phases regardless of ongoing behavioral performance, as long as mice retained consistent or escalating daily infusions. All maintenance sessions followed an intermittent access (IntA), FR1 schedule. IntA sessions were 5 hours and 5 minutes long and consisted of 10 repeating, 30-minute-long blocks. Mice began with a 5-minute baseline recording in the self-administration box before levers and cue lights were presented to them. Each block began with a 6-minute FR1 period in which all active lever presses were rewarded with intravenous fentanyl delivery followed by only a 3-second timeout (the duration it takes to infuse the drug). Then, while levers remained accessible, cue lights were turned off during the subsequent 24 minutes, and active or inactive lever presses had no consequences but were recorded. This block would repeat 10 times to provide a total of 1 hour of access to fentanyl per day, over a 5-hour session.

### Video recording

All behaviors were video-recorded using two NeuroMotive (Blackrock Microsystems, Inc., Salt Lake City, UT) camera systems for a top- and side-view of the arena. Cameras were synchronized with one another and with the neurophysiological data acquisition system. For locomotor-related behaviors, top-view videos were tracked using Ethovision X12 (Noldus, Wageningen, the Netherlands) and changes in center-point location were used to determine the distance traveled.

### Withdrawal severity index

Behavioral quantification of opioid withdrawal severity has traditionally relied on summing the number of observed behaviors or applying arbitrary weights to individual behaviors^46,58–62^. To develop an unbiased, data-driven measure of withdrawal severity, an experimenter blinded to condition manually scored multiple behavioral indices associated with opioid withdrawal, including defecation, forepaw tremors, rearing, grooming, jumping, booty dragging (walking with rear tucked under, dragging along the floor), and orofacial behaviors (swallowing, gaping, and teeth chattering). In addition, immobility was quantified as the cumulative duration of bouts >2 s during which the center point of the mouse moved <1 cm per second.

Each behavioral variable was independently *z*-scored across all mice within a cohort of saline-treated and morphine-withdrawing animals to account for inter-animal variability. The resulting *z*-scored behavioral matrix was submitted to principal component analysis (PCA; **Fig. S6J**). The first principal component (PC1) explained ∼54.2% of the total variance and effectively separated withdrawing from non-withdrawing mice (*p*<0.0001; Welch’s *t*-test; **Figure S6K**). PC1 loadings revealed a natural weighting structure in which certain behaviors (e.g., immobility, orofacial behaviors, jumping, forepaw tremors, defecation, and booty dragging) contributed positively to withdrawal severity, whereas others (e.g., grooming and rearing) contributed negatively (**Fig. S6L**). PC1 scores for each mouse were therefore used as a single composite withdrawal severity index reflecting the relative severity of behavioral withdrawal. These data were derived from the escalating morphine withdrawal cohort.

### Neural electrophysiology data acquisition

Neurophysiology data were acquired using a Cerebus acquisition system (Blackrock Microsystems, Inc., Salt Lake City, UT). Animals were connected to the system using a Mu-32 channel headstage (Blackrock Microsystems, Inc., Salt Lake City, UT), which was attached to a motorized HDMI commutator (Doric Lenses, Quebec, Canada). Local field potentials (LFPs) were bandpass filtered at 0.5-250Hz and sampled/stored at 1kHz. All LFP data were referenced to a ground wire connecting the ground screws above cerebellum and anterior cranium.

### Cellular unit acquisition and isolation

Online visual and auditory inspection of each channel was used to determine if unit activity could be isolated. Channels for which single and multi-units were suspected were referenced against a channel from the same brain area that did not have any clear units. If no wires within the same region were available, the next closest brain region was used to locally reference the channel. Neural activity was high pass filtered at 250 Hz and sampled at 30 kHz. Only activity that crossed a predetermined threshold (approximate signal-to-noise ratio > 3:1) was stored. After data collection, cells were sorted offline using the Plexon Offline Sorter (Version 4.6.2, Plexon Inc., TX). Cell clusters that were clearly isolated from background noise were saved for final analysis. Clusters were defined by visual inspection from a well-trained experimenter based on waveform properties such as peaks, valleys, non-linearity, and software-generated principal components. Single and multi-units were used for analysis.

### LFP processing to remove signal artifact

Our prior heuristic approach to eliminate recording segments containing non-physiological signals^26,63–65^ was adapted and utilized as described here. We first computed the signal envelope for each channel by utilizing the magnitude of the Hilbert transform. For any 5-second window in which the envelope surpasses a predetermined low threshold, we discard the entire segment if, at any point within that window, the envelope exceeded a second, higher threshold. The two thresholds were independently determined for each brain region. The high threshold was set to 25 times the median absolute deviation of the envelope value specific to that region. We chose such a high threshold since fentanyl and morphine injections elicit a very strong neurophysiological response that exceeds multiple deviations more than non-drug windows. We found that this threshold protected against removal of windows with strong responses to the drugs while also remaining sensitive enough to remove ‘noisy’ time windows based on visually inspected LFP power plots. The low threshold was empirically established as 3.33% of the high threshold. If more than half of a given 5-second window was removed for any given channel, we removed that entire window for all channels. Additionally, all windows in which the standard deviation of the channel was less than 0.01 were excluded. This resulted in an average of 0.6% ± 1.9% of windows per mouse being removed. Then, 60-Hz line artifacts were removed using a series of Butterworth bandpass filters at 60 Hz and harmonics up to 240 Hz with a stopband width of 5 Hz and a stopband attenuation of -60 dB (buttord; MATLAB R2021a). Finally, signals were downsampled to 500 Hz for feature calculations. All processing steps described here were done using MATLAB R2021a (The MathWorks, Inc., Natick, MA, USA).

### Feature extraction

Feature extraction was performed identically to previous works^26,63–65^. Briefly, LFPs were averaged across wires within the same region to generate a composite LFP measure. Signal processing was conducted using MATLAB (The MathWorks, Inc., Natick, MA). For LFP power, Welch’s method was applied to the averaged LFP signal using 250-sample rectangular sub-windows (0.5 seconds at 500 Hz) with 100-sample overlap (0.2-second step). Power spectral density was estimated from 1–56 Hz in 1-Hz bins. LFP cross-regional coherence was computed from pairs of averaged LFP signals using Welch’s magnitude-squared coherence with 0.5-second rectangular sub-windows and 0.2-second overlap. Coherence was estimated as a function of the auto-spectral densities of regions A and B (|*Psd*_*AA*_| *=* |*Psd*_*BB*_|) and the magnitude of their cross-spectral density : (|*Psd*_*AB*_|*=* |*Psd*_*BA*_|)

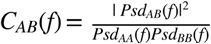

To capture dynamic neural responses to repeated fentanyl exposure, we computed LFP spectral power and coherence features in 5-second windows for each recording session.

### Tensor component analysis (TCA)

#### Input tensor

Each fentanyl dosing session began with a 2-minute baseline home cage recording before animals were placed into a monitoring box for at least 5 minutes to acclimate and then injected with saline or fentanyl. To account for differences in scale between power and coherence features, feature values were normalized within each feature class (power or coherence) prior to tensor construction. For power features, we computed log-transformed values as *log(power + ε)*, where *ε* = 10 ^12^ X *median(power)*, averaged values within baseline (first 2 minutes) and post-injection (1–5 minutes after injection) periods, defined as the post-pre difference (log fold-change). For coherence features, values were clipped to [IO ^−6^, 1−10^−6^], transformed using the logit function 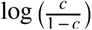, averaged within baseline and post-injection periods, and expressed as a post-pre difference (log-odds change).

#### Regularized canonical polyadic (CP) decomposition

To uncover a low-dimensional structure within the data, we applied an unsupervised canonical polyadic (CP) tensor decomposition that approximates the multidimensional tensor into a sum of ***R*** rank-one tensors for a user-specified ***R*** . The tensor denoted by *X* ∈ ℝ^*I*×*J*×*K*^, which consists of *I* mice, *J* baseline-subtracted LFP features, and *K* experimental days, was masked with NaNs to ignore missing values (e.g. recording days with poor signal quality) and to preserve tensor structure.

Our custom CP decomposition model approximated the data tensor as a sum of *R* rank-one components. The model was parameterized by three low-rank factor matrices: *A* ∈ ℝ^*I*×*R*^ for mouse loadings, *B*∈ ℝ^*J*×*R*^ for LFP feature loadings, and *C* ∈ ℝ^*K*×*R*^ (day factor loadings), where *R*is the user-specified number of components. We use “component” to refer to one rank-one term in the decomposition, indexed by *r* . Each component *r*, is parameterized by corresponding loading vectors (*A*_:,*r*_, *B*_:,*r*_, *C*_:,*r*_), and their outer product *A*_:,*r*_*°B*_:,*r*_*°C*_:,*r*_ defines a rank-one component, where o denotes the outer product between vectors. Thus, *B*_:,*r*_ is the LFP feature-loading vector for component *r*, and it can be visualized as a candidate network-level pattern because it describes which LFP features covary within that component. In this notation, : denotes all entries along a given dimension. A range such as *start:finish* selects only a contiguous subset along that dimension. For example, *B*_*start:finish, r*_ would select only the LFP features indexed from start through finish for the same component *r* . This low-rank formulation, analogous to PCA, represents the data as a small set of latent components that capture patterns shared across mice, features, and days, such that:

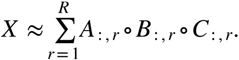

The decomposition was optimized using gradient descent implemented in PyTorch to minimize the sum of squared errors (SSE).

#### Rank selection

To determine the appropriate model complexity, we performed a rank sweep over *R= {1–50}*. For each rank, 10 models were fit with different random initializations. For each of the 10 replicate fits, an averaged Z-scored SSE and averaged Z-scored cosine distance were summed together for a *composite score*. The minimum composite score across all tested *R* was chosen as the optimal number of total ranks, *R*, that balanced model complexity and rank reproducibility.

#### Stability-based component discovery

Using the selected number of total ranks, *R*, we performed 1000 replicate model fits using varying random seeds and storing factor matrices for each replicate. Each replicate consisted of *R* components.

To align LFP feature loading vectors across replicates, we computed similarities between L2-normalized LFP feature loadings (*B*), yielding an *R×R* similarity matrix for each pair of models by using the following procedure: Each LFP feature loading vector was normalized prior to computing absolute cosine similarity to ensure that component matching was based on similarity in feature patterns independent of scaling. Absolute cosine similarity was used to account for sign indeterminacy. We used the Hungarian algorithm to determine the optimal one-to-one correspondence of components across replicate models, maximizing total similarity across matched pairs and resolving permutation indeterminacy in component ordering. We then calculated average cosine similarity between each column vector in *B* corresponding to a single component.

To capture stable LFP feature loading vectors, we retained vectors that were consistently matched across at least 90% of replicates and had a mean similarity of ≥0.90. These retained loading vectors were then clustered using absolute cosine similarity of concatenated corresponding normalized *A*_:,*r*_, *B*_:,*r*_, and *C*_:,*r*_, vectors for all *r* to look for complete components, rather than matching on *B*_:,*r*_, alone. Within each cluster, the model replicate with the highest average similarity to other members was selected as the representative centroid. Finally, centroid components were deduplicated by comparing each pair under the four sign-flip patterns consistent with CP’s inherent sign ambiguity (flipping the sign of any two of the three loading vectors), collapsing pairs whose corresponding *A*_:,*r*_, *B*_:,*r*_, and *C*_:,*r*_, loading vectors matched above threshold. The retained loading vectors for each surviving component served as its final representative *A*_:,*r*_, *B*_:,*r*_, and *C*_:,*r*_.

Final factors were aligned so that the average LFP feature loadings were positively oriented. This alignment ensures consistency across components and enhances interpretability by standardizing the directionality of feature weights across replicate fits, facilitating downstream comparisons.

#### Training and testing

To evaluate whether the final components generalize to unseen animals, after fitting the TCA on the training data, we projected the holdout data onto the learned LFP feature vector, *B*_:,*r*_, for each retained component. Model training incorporated early stopping based on tolerance and patience criteria. This formulation ensures that the learned latent structure reflects both biologically interpretable constraints and statistical sparsity. Then, we computed a feature-space projection that quantified the engagement of a given component within the new data. For a given holdout tensor *X*_*holdout*_ ∈ ℝ^*I*×*J*×*R*^, the engagement of the *r*^*th*^ component for mouse ***i’*** on day ***k*** was computed as the dot product between the observed feature vector and the LFP feature loadings corresponding to the *r*^*th*^ component from the training model:

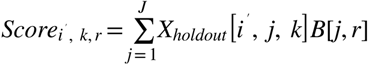

This approximate projection assumes near-orthogonal LFP feature loading vectors across components and yields a matrix of component engagement values of size ***I’ *K*** for each component ***r***, reflecting how strongly the feature pattern associated with that component is expressed for each mouse across days. Because the projection is performed directly in feature space, the resulting engagement values capture the joint expression of the mouse and day components without requiring additional optimization or tensor decomposition on the holdout data. This approach preserves the feature structure learned from the training cohort while providing a simple and interpretable measure of component activity in independent animals.

### Discriminative Cross-Spectral Factor Analysis – Nonnegative Matrix Factorization (dCSFA-NMF)

#### Label assignment for training dataset

To train and calculate predictive performance within a supervised classification model, binary state labels (withdrawal vs. non-withdrawal) must be defined for all windows across datasets. To prevent ambiguous labeling, we chose to take time points in which we were most certain mice were in or not in withdrawal for training. A total of 21 mice (24,256 5-second windows) were used to train the model. 14 mice (8 male, 6 female) received chronic fentanyl-naloxone (FN), 7 mice (4 male, 3 female) received chronic saline injections, and 4 of the 7 chronic saline mice were reused for acute fentanyl-naloxone (AFN) recordings (2 male, 2 female). For all mice, day 1 recordings in the home cage and the first 15 minutes in the monitoring box were used as negative-class (0) non-withdrawal windows. For chronic saline and AFN mice, the last 60 minutes of day 7, the period following the last injection of saline or naloxone, were additionally used as negative-class, non-withdrawal data. The same 60-minute window for FN mice, following naloxone precipitation, was taken as positive-class (1) withdrawal windows. AFN mice were included as controls to ensure that the network does not simply learn a negative-valenced experience, as acute fentanyl-naloxone is known to induce a highly aversive experience for individuals. Additionally, to ensure that the model does not simply learn behaviors exclusive to withdrawal (e.g. jumping) or changes in locomotion, we chose to calculate neural features using relatively long windows (5-second windows) for an entire 1-hour period that encompasses a wide variety of behaviors. This also encourages the model to learn a withdrawal state independent of behaviors that may or may not be present at a moment-to-moment basis. FN mice that did not jump were removed from the training dataset.

#### Label assignment for testing datasets

Model performance was assessed in two independent holdout datasets consisting of new cohorts of mice withdrawing from morphine. For constant morphine dosing, 10 mice (3 male, 7 female) were treated with morphine, and 6 (3 male and 3 female) with saline. Performance was assessed on the last day of dosing. All data for saline-treated mice were classified as negative-class, non-withdrawal data. For morphine-treated mice, the initial 27 minutes of neural recordings (in which mice exhibit spontaneous withdrawal) and the last 60 minutes (following naloxone-precipitated withdrawal) were labeled as positive-class, withdrawal data, whereas the 150-minute period following morphine (which suppresses withdrawal) was labeled as negative-class, non-withdrawal data (**Figure 3A**).

For escalating morphine withdrawal, recordings took place 12–16 hours after the final saline or morphine injection. Here, all data from saline-treated mice were labeled as negative-class, non-withdrawal data and all data from morphine-treated mice were labeled as positive-class, withdrawal data.

#### Training, validation, and test splits

Mice were split into three groups for model hyperparameter tuning, training, performance assessment: training data, validation data, and testing data. These splits were performed by mouse, such that all data belonging to a mouse were contained in a single group and not used across groups. Splitting by mouse is critical as it prevents a machine learning model from simply learning the identity of a mouse in the training data to achieve inflated performance on holdout data from the same mice. Additionally, we wished to see how our model performs on data from completely new subjects, which is a situation analogous to the conditions of a clinical setting. Training and validation data were used for model development, where many sets of hyperparameters and model formulation were tested and optimized for prediction and reconstruction of the data. Test data were kept as true holdout data, upon which we neither observed nor tested our model until the final model architecture was determined. Once hyperparameters were determined, the training and validation data were combined for one final training dataset, and predictive performance was assessed on the test data.

#### dCSFA-NMF model description and fit

Discriminative Cross-Spectral Factor Analysis – Nonnegative Matrix Factorization (dCSFA-NMF) is a machine learning framework for discovering predictive factors relevant to a behavioral assay or emotional state of interest^66,67^. This method has been used previously to detect brain networks corresponding to stress, social activity, anxiety, and aggression in mice using LFP data^26,63–65^. To clarify terminology, we use “factor” in dCSFA-NMF to refer to a latent model component, not to the loading vector itself. This differs from the CP/TCA terminology above, where the decomposition is described in terms of matrices (***A***,***B***, and ***C***) and a component is defined jointly by one loading vector from each matrix. Each dCSFA-NMF factor/component has an associated LFP feature-loading vector in ***W*** and an activity score in ***S***, defined below. When the feature loadings form a coherent pattern across power and coherence features, we interpret that latent factor/component as a candidate network.

Like other factor models that are used in neuroscience, such as PCA, independent components analysis (ICA), and NMF, dCSFA-NMF identifies underlying components interpreted to be networks of connectivity. The superposition of these networks then explains the observed neural activity. While the previously mentioned unsupervised methods identify networks that maximize explained variance, dCSFA-NMF aims to discover networks that balance how well networks predict a target of interest with how much variance they explain. Because emotionally relevant brain states may not explain a large fraction of the variance in the LFP data, unsupervised methods can miss relevant brain networks. To compensate, dCSFA-NMF makes use of an additional supervised term, where the activity score, or network expression, of one or more supervised factors is correlated with a behavior or emotional state of interest.

Specifically, the model learns *K* fixed factor loading vectors that are concatenated together into a matrix *W* ∈ *R*^*K*\**M*^. Each factor loading vector *W*_[:,*k*]_ represents the usage of different LFP features and can represent a “network.” These networks can reconstruct observed data *X* ∈ *R*^*N*\**M*^ using an array of network activity scores *S* ∈ *R*^*NK*^ such that *X* = *SW. W* and *S* are also constrained to be positive as the features of use - power and coherence - are also nonnegative. Network activity scores are inferred from the observed data using an encoder function *S*_[*n*,:]_ = *f*_*θ*_(*X*_[*n*,:]_), which takes the form of a variational autoencoder. For *Q* ≥ 1 supervised networks, the activity scores for each subject *n, S*_[*n*, 1:*Q*]_ ∈ *R*^*Q*^, are used in a logistic regression model *fΦ* to predict the behavior of interest *Prob*(*y*_*n*_ = 1) = *f*_*Φ*_(*S*_[*n*, 1:*Q*]_). This formulation explicitly constrains our predictions to use a sparse combination of all networks, namely only the supervised network(s), to narrow the scope of our network discovery and reduce the total number of comparisons in downstream testing. The parameters of the model are then optimized using the loss function:

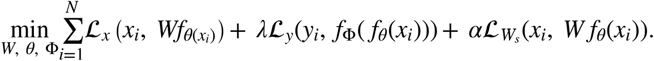

Here, *L*_x_ is the reconstruction error between the original power and coherence features and those generated by the product of our network scores and networks, ***sW***. In this work we make use of the Mean-Squared-Error (MSE) function. Our predictive loss *L*_y_ is a binary cross-entropy loss and penalizes our model for incorrectly predicting the behavioral state of each window. The impact of the predictive loss can be tuned using the hyperparameter *λ*. Lastly, we impose a second reconstruction loss, 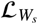, on the supervised network scores.

This reconstruction loss prevents our neural network encoder ***f***_*θ*_ from learning an uninterpretable near-zero noise embedding for the supervised scores that predicts the withdrawal vs. non-withdrawal state well, but has little to no contribution to explaining neural activity. This loss can be formulated as another MSE loss between the outer product of the supervised network activations and the supervised electome network and the features. Alternatively, this loss can be formulated as a penalty to drive the supervised network scores to reconstruct the residual of the unsupervised networks and features. We used the latter in our analysis.

We have previously found that pretraining our model factor loading vectors and encoder weights provided a substantial improvement on representational stability and predictive accuracy^64^. We performed pretraining by training a traditional NMF model on our data and then sorting the factor loadings based on their correlation with the behavior of interest, so that more correlated networks would be selected for supervised use. We temporarily froze the weights of our sorted NMF factor loadings, then trained the encoder (***f***_*θ*_) parameter to estimate network activity scores and finally trained the classifier (*f*_*Φ*_) parameters to predict corresponding labels based on those scores. We found that training the encoder for 300 epochs and training the classifier for 10000 epochs was sufficient for the optimization algorithm to converge and stabilize at a minimum of the loss function. This large number of epochs for the classifier was predominantly due to the long training time it took to converge and stabilize the loss function when ***K*** = 1. After pretraining, we then unfroze all parameters and trained them jointly. We found that an additional 200 epochs were sufficient for the training to converge and stabilize at a minimum of the loss function.

#### Hyperparameter selection strategy

The dCSFA-NMF model requires selection of several hyperparameters. These hyperparameters include the number of electome networks *K*, the importance of the supervised task 2, and the importance of the supervised network reconstruction *α*.*Q*, the number of supervised networks, was set to 1 for simplicity. Generally, the number of electome networks controls how well we can reconstruct the original LFP data but does not greatly affect the overall prediction quality. Additionally, we constrained our model to only identify supervised networks with scores that positively correlated with predicting a heightened withdrawal state, as we aimed to discover neural features activated by withdrawal rather than those inhibited by withdrawal.

We chose the hyperparameters in a sequential process. We first chose hyperparameters to optimize predictions of withdrawal vs. non-withdrawal states, which in our prior experience had little dependence on *K*. Thus, for this stage we used a nominal value of 6 for *K*, whose value will be more carefully selected in the next stage of the process. Models were iteratively trained on train–validation splits in which 4 mice (2 saline and 2 FN) of the 21 total mice were randomly selected for validation and training proceeded on the remaining 17 subjects. We tested various hyperparameters relating to reconstruction weight, supervision weight, reconstruction weight for the supervised network specifically, the type of optimizer used (Adam or stochastic gradient descent), learning rate, and the number of epochs used for training, pretraining the encoder, and pretraining the decoder. The set of hyperparameters that optimized predictions of withdrawal vs. non-withdrawal states in the train–validation splits were used going forward.

To choose the value of *K* in the next stage, we performed a grid-search cross-validation using *K*= {1, 2, 3, … 20} with the optimized hyperparameters from above. For each value of *K*, a random seed (reproducible random initialization) was selected, and 10 separate models were trained. *R*^2^ (proportion of variance explained by the model’s reconstruction of the original LFP data), area under the receiver operating curve (AUC; metric to evaluate predictive performance of withdrawal vs. non-withdrawal labels), and cosine similarity values (metric of network consistency) were calculated across all 10 folds (**Figure S3**). Given that each of these metrics are bound between [0,1], we calculated a simple “combined score” by multiplying the average value of each of the three metrics together such that a model with a combined score of 1 would have perfect reconstruction, prediction, and model stability. This led to a model with *K*=5; one supervised network primarily reflecting the withdrawal state and 4 unsupervised networks that likely reflect other neural processes.

#### Performance metrics – predictive modeling

To evaluate the predictive performance of our model, we use the area under the receiver operating curve (AUC), as mentioned briefly above. This metric is common in machine learning literature and can be viewed as a class rebalanced accuracy between [0,1], where an AUC of 0.5 indicates chance performance for binary classification, AUC of 1 indicates perfect prediction, and an AUC of 0 indicates that the model is predicting the exact opposite class assignment, such that all negative-class data is predicted as positive-class, and vice versa. AUC significance can be evaluated using the Mann–Whitney U statistical test compared to chance performance (i.e., AUC=0.5 for binary predictions).

For evaluating our models, we obtain an AUC for each mouse and then report the group mean and standard error of the mean (SEM) for each paradigm when possible. As class assignment is not always perfectly matched between the number of positive- and negative-class windows, reporting of the AUC per mouse addresses the possibility of our model overfitting to the neural activity of mice in paradigms with imbalanced or increased number of samples/windows. Furthermore, emotional states such as withdrawal are complex and have heterogeneous presentations across individuals. By reporting AUC by mouse, we allow opportunities for post-hoc analyses of data from mice with heterogeneous presentations.

Given that mice treated with only saline or those in experiments in which neural activity is only collected during the withdrawal phase (as is the case for the escalating withdrawal experiments), these mice will only contain a single class of labels (withdrawal or non-withdrawal). As such, per-mouse AUCs cannot be calculated for these mice. When this occurs, average network activation per mouse is calculated, grouped per condition, and the mean and standard deviation between groups are compared using appropriate statistics.

#### Performance metrics – generative modeling

We evaluated how well our models explain neural activity by taking the MSE of the model’s predicted power and coherence features compared to the originally observed values.

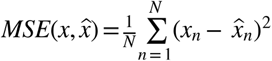

During model training, we weighted the reconstruction of each of our feature types (power and coherence) by their prevalence, such that power holds equal importance to coherence despite representing a smaller number of features.

In holdout datasets, we report the reconstruction performance using the coefficient of determination ***R***^***2***^ per mouse as follows:

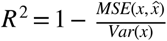

#### Performance metrics – model consistency

To evaluate consistency of model architecture, we used the cosine similarity formula which calculates the angular distance between two vectors on a scale of [0,1]. Values range from 0 (orthogonal vectors) to 1 (perfect alignment), as all dCSFA-NMF factor loadings (***W*** ) in this scenario were constrained to be nonnegative. For concreteness, when evaluating the cosine similarity between two networks found by dCSFA-NMF, we have their representations *W*[:,*k*] and *W*[:,*j*] Their cosine similarity is given by the following formula, where “•” denotes the dot product and || • ||_2_ denotes Euclidean (L2) norm:

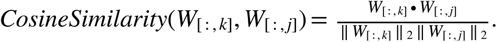

#### Single feature-group models

To assess the necessity of the full network of regions compared to its constituent regions alone or in pairs, we trained independent dCSFA-NMF models on all single regions or region-pairs, each referred to as a single feature-group model. Single-region models contain only power features from 1–56 Hz for that given region. For region-pair models, power within each of the two regions, as well as coherence between regions, all within frequencies ranging from 1–56 Hz, were used to train the models. Models were trained with network-optimized parameters on the same data used to train the full network. Each model was then used to assess if any single feature-group could predict withdrawal in holdout morphine datasets.

#### Statistical analysis

Data were analyzed using GraphPad Prism 10.5.0 and Python (SciPy; statsmodels). Results are reported as mean +/-SEM (error bars or shaded areas). Normality was formally assessed using the Shapiro–Wilk test for all comparisons besides the single feature-group models (**Figure S5**). Normally distributed data were analyzed using parametric tests and non-normally distributed data with corresponding non-parametric alternatives. Statistical significance was set at α=0.05 and conservative Bonferroni corrections were used any time multiple hypothesis testing was performed. For multi-factor analysis, we used mixed-effects models when data contained both between- and within-subject factors (or when repeated measures were unbalanced), and two-way Repeated Measures ANOVAs when all factors were within-subject and normally distributed. If main or interaction effects *reached p*<0.05, we performed post hoc tests (pairwise t-tests, Dunnett’s test, and Sidak’s test where appropriate) with Bonferroni corrections. Unpaired (Welch’s) or paired t-tests were used for single comparisons between normally distributed groups to avoid any assumptions of equal variance between groups, whereas Mann–Whitney U or Wilcoxon signed-rank tests were used for non-normal data. Pearson or Spearman tests were applied to assess correlations between normal or non-normal variables, respectively. All analyses used two-tailed statistics, except where specified.

Our a priori hypothesis was that morphine-withdrawing mice would show elevated network activity relative to saline controls across the home cage, spontaneous withdrawal, and naloxone-precipitated withdrawal contexts. These three phases were recorded continuously within a single test-day session in the same animals; thus, we modeled Group (morphine vs. saline) as a between-subject factor and Time (context) as a within-subject factor in a two-way RM-ANOVA, testing whether the predicted group difference held across all three contexts.

Network activity following the subsequent morphine injection was assessed separately, with a two-tailed Welch’s t-test comparing morphine- and saline-treated mice at that timepoint. This addressed a different question than the RM-ANOVA: not whether the elevation generalized across contexts, but whether it could be pharmacologically reversed. Here we predicted the opposite pattern: morphine rescue would abolish the group difference.

We then used two additional two-tailed Welch’s t-tests to evaluate network activity 24 hours and 5 days after the rescue injection, in independent recording sessions separate from the test-day protocol above. We predicted the group difference would still be present at 24 hours but not at 5 days based on the established time courses of opioid withdrawal in mice^3^.

When evaluating the ability of single feature-group models to predict withdrawal in holdout datasets, data from both morphine withdrawal experiments (constant and escalating) were combined to increase sample size. Because this approach introduced a mixture of paired and unpaired observations across experiments, we used a linear mixed-effects model with mouse identity included as a random intercept to account for repeated measures within subjects when comparing group differences.

Statistical significance for unit–network coupling was assessed using a nonparametric permutation test. Spike times (or unit identities) were randomly shuffled relative to network activity across 1,000 iterations to generate a null distribution of coupling values, against which the observed coupling was compared. This approach avoids assumptions of normality and accounts for the temporal structure of the data. For the conditioned place aversion experiment, we employed a biased design in which mice always underwent fentanyl–naloxone reversal in their initially preferred chamber. A conditioned aversive response was defined as a significant decrease in time spent in the drug-paired chamber post-conditioning relative to pre-conditioning, along with a significantly lower preference than chance (50%). This statistical criterion mitigates a common limitation of biased designs; namely, that changes in preference could otherwise reflect chamber habituation rather than true conditioned aversion. No statistical methods were used to predetermine sample size. Mouse cages were pseudorandomly assigned to treatment or control conditions. Experimenters were not blinded during data collection or analysis, except for blinding the experimenter during manual behavioral scoring of withdrawal ethograms.

#### Visualization

All plots were made using GraphPad Prism 10.5.0 or Python (Matplotlib and Plotly packages) and subsequently edited for visualization and aesthetic purposes only in Adobe Illustrator 28.0 when necessary. All schematics were made using Adobe Illustrator 28.0. Circos plots were visualized using custom Python code. All figures were constructed using Adobe Illustrator 28.0.

#### Reproducibility

Preprocessing and feature extraction code was performed in MATLAB (MathWorks) R2022a using the LFP feature extraction pipeline found here: https://github.com/carlson-lab/lpne-data-analysis. A PyTorch implementation of dCSFA-NMF can be found at https://github.com/carlson-lab/lpne/. All code for TCA and dCSFA-NMF network generation, hyperparameter tuning, model implementation, and plotting for replicating our results is available at https://github.com/carlson-lab/FentanylWithdrawal and will be made publicly accessible upon publication.

## ACKNOWLEDGEMENTS

We would like to thank Jean Mary Zarate for comments on this work; Morgan Gallimore for technical support; and Minel Arinel for substantial support and comments on figure design.

## AUTHOR CONTRIBUTIONS

Conceptualization and Methodology – KA, KKW-C, SDM, and KD; Formal Analysis – KA and KMW; Investigation – KA, KKW-C, CB, KMW, and MAG; Resources – SDM and KD; Writing – Original Draft – KA; Writing – Review & Editing – KA, KMW, KKW-C, SDM, DEC, and KD; Visualization – KA; Supervision – KA, DEC, and KD; Project Administration and Funding Acquisition – KA and KD.

