## Supplemental Materials for "A widespread internal brain state for fentanyl withdrawal"

### SUPPLEMENTARY TABLE S1: DETAILED AUTHOR CONTRIBUTIONS

| Author | Contribution |
|---|---|
| **Karim Abdelaal** | Conceived of and led the withdrawal electome project and all experiments within the paper; designed all experiments within the paper; built electrodes with CB; implanted mice for all data in the paper; analyzed all behavioral data with KW; processed and analyzed all neurophysiology data and machine learning analysis, including model discovery, validation, projections, and statistical testing; analyzed unit-network coupling data; jointly performed histological confirmations with MAG; supervised all data collection and analysis; wrote the paper and created all visualizations with edits from SDM, JZ, and KD. |
| **Kathryn K. Walder-Christensen** | Assisted in data collection for fentanyl withdrawal, constant morphine, and escalating morphine experiments with KA and CB; assisted in electrode construction for both morphine experiments; jointly piloted withdrawal experiments with KA; assisted in experimental design and conception of fentanyl and morphine withdrawal experiments with KA. |
| **Cameron Blount** | Built electrodes with KA; ran fentanyl withdrawal behavioral experiments with KA. |
| **Kellie M. Williford** | Quantified all behavioral ethograms for all withdrawal severity measures; processed and isolated all unit firing data from fentanyl and morphine cohorts; contributed to writing methods sections of the manuscript; provided feedback and edits on the manuscript. |
| **Marisella Adams-Grimaldi** | Jointly performed all histological confirmation, including slicing, mounting, and imaging with KA. Assisted in electrode construction with KA. |
| **Stephen D. Mague** | Assisted in project design and methodological considerations. Revised the manuscript. |
| **David E. Carlson** | Oversaw all machine learning based analyses with KD. Revised the manuscript |
| **Kafui Dzirasa** | Supervised and oversaw all experiments and analyses with DEC. Secured resources. Revised the manuscript. |

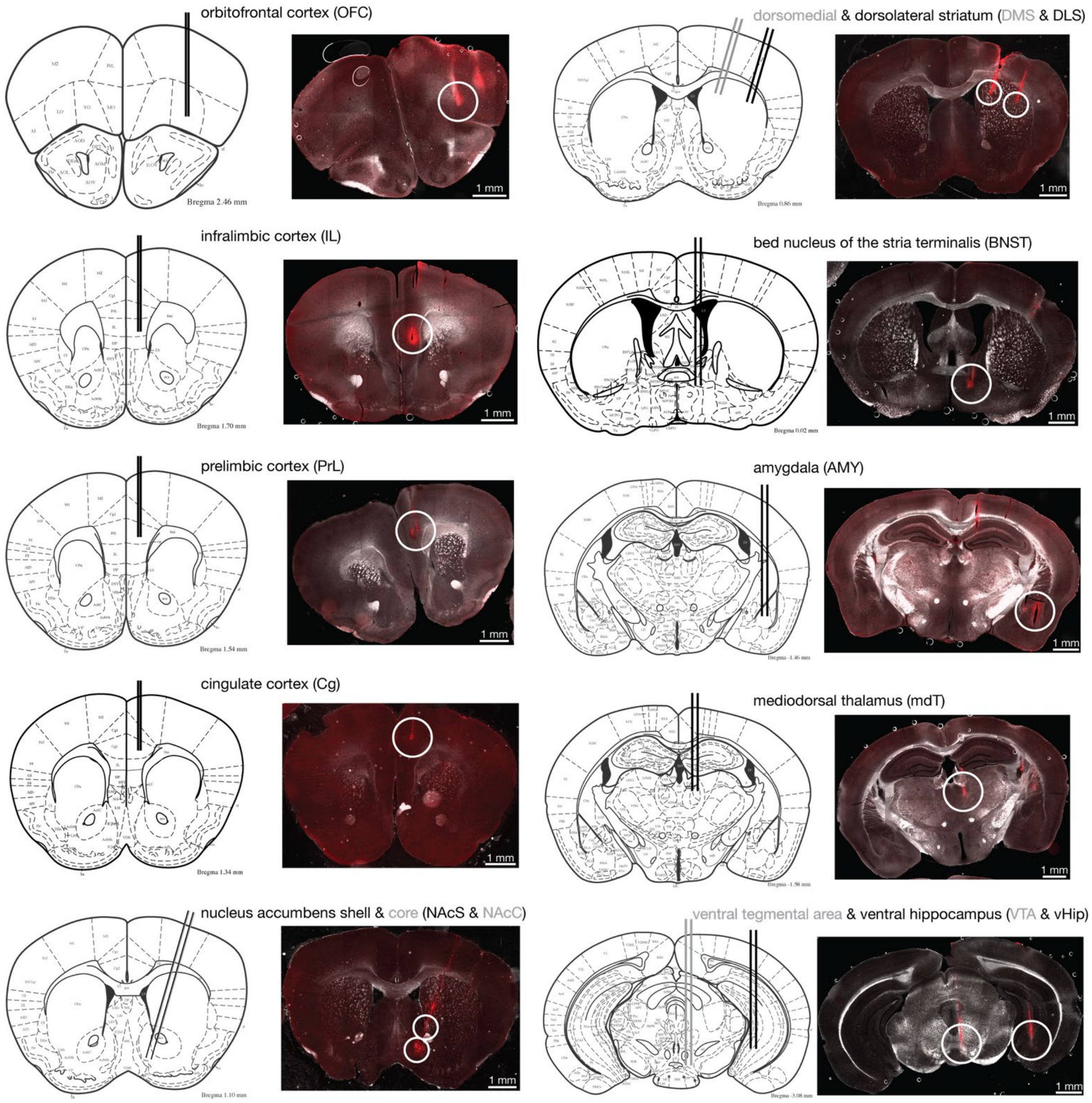


**Figure S1: Electrophysiological target verification.** Representative electrode placements for all recording targets. Electrode tracks were visualized in thin post-mortem sections using brightfield imaging to identify anatomical landmarks and red-filtered fluorescence to detect dye-coated electrode tracks. Aside from these regions, all mice were also implanted with electrodes targeting the anterior insula, central amygdala, and periaqueductal gray; however, due to inconsistent targeting or inability to confirm placement in some animals, these regions were excluded from all analyses for all mice. A total of 121 mice were implanted for this study, 11 were removed for inaccurate electrode placement, and an additional three were removed due to an inability to confirm electrode placement from histological samples.

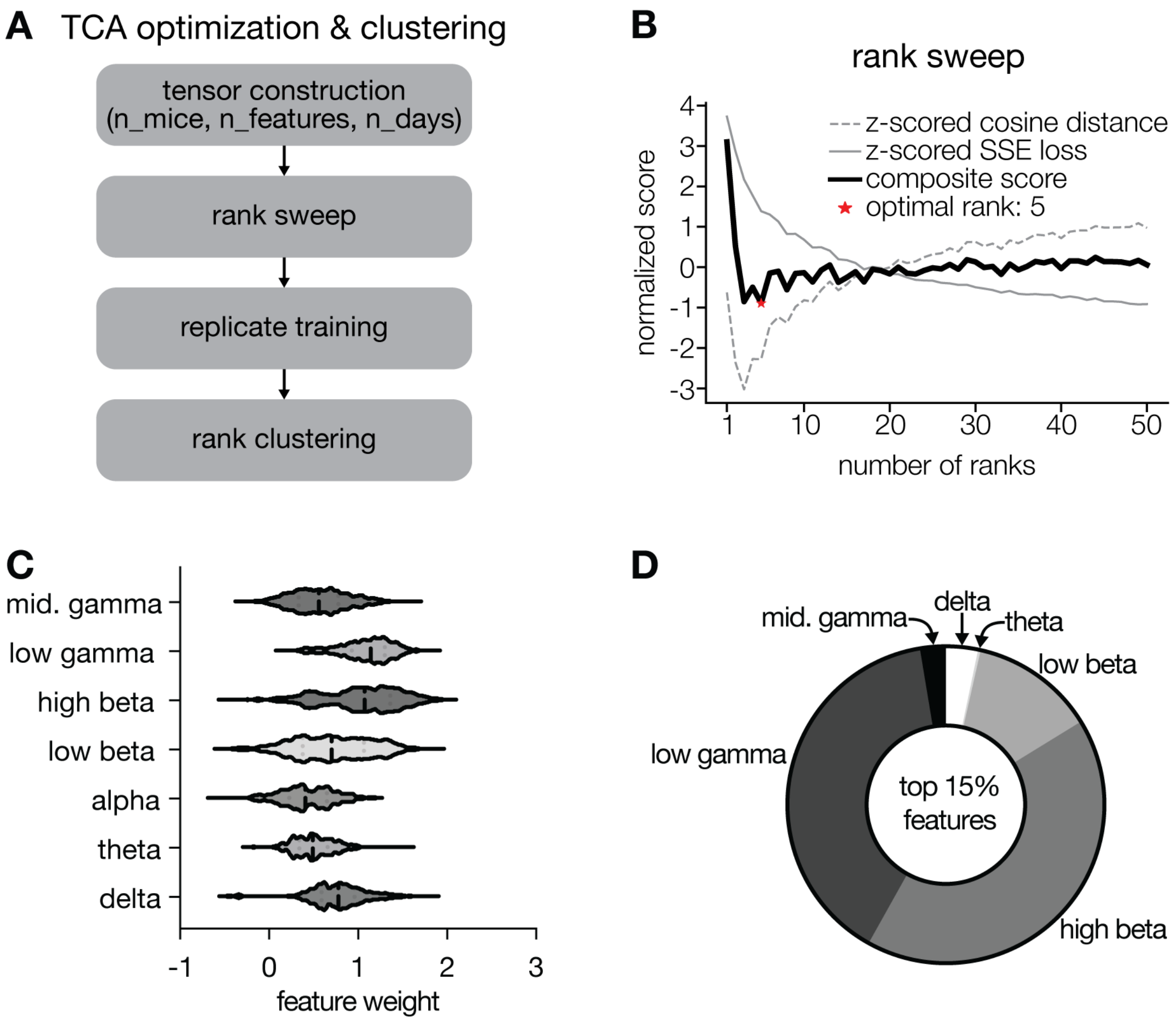


**Figure S2: Tensor component analysis (TCA) optimization steps.** (A) Data were first organized into a three-dimensional tensor (mice × features × dosing days) and trained across *k* ranks, from *k*=1–50, 10 times each. This rank sweep provided the optimal number of ranks per model. Then, replicate training of models with random starting seeds were repeated 1,000 times and clustered to identify highly stable and recurring components across replicates (see Methods for details). (B) Rank sweep showing model performance across candidate ranks, with an optimal rank of 5. Optimization was defined as the minimum value of the summed z-scored cosine distance and z-scored sum-of-squares error (SSE) loss. (C) Feature weights across frequency bands. Activity is highest in the high beta (20–30 Hz) band, followed closely by low gamma (30–40Hz) activity. Activity in alpha band is lowest. (D) Number of frequency-band specific features within the top 15% of model weights. High beta is most represented, followed closely by low gamma activity.

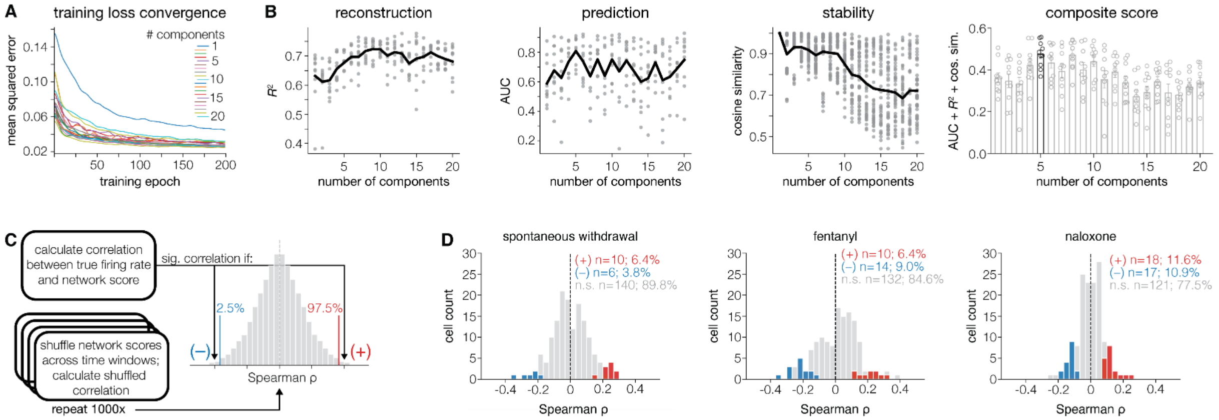


**Figure S3: dCSFA-NMF model optimization and validation of unit–network coupling.** (**A**) Mean-squared error across training epochs for models with 1–20 total networks shows that the training loss function has stabilized by ~200 epochs, implying learning has completed. (**B**) The optimal number of networks was determined by jointly maximizing the average reconstructive accuracy ($R^2$), predictive performance (area under the receiver operating curve; AUC), and network stability (cosine similarity) across 10 models trained with varying numbers of total components. This yielded a final dCSFA-NMF model with five total networks. **(C)** Schematic of the permutation-based approach used to assess unit–network coupling. For each unit, the Spearman correlation between firing rate and network score across time was compared to a null distribution generated by 1,000 circular shuffles of the network time series; units >97.5$^{th}$ or <2.5$^{th}$ percentile were considered significantly positively or negatively coupled, respectively. **(D)** Distribution of positively, negatively, and non-correlated units across spontaneous withdrawal, fentanyl, and naloxone time periods.

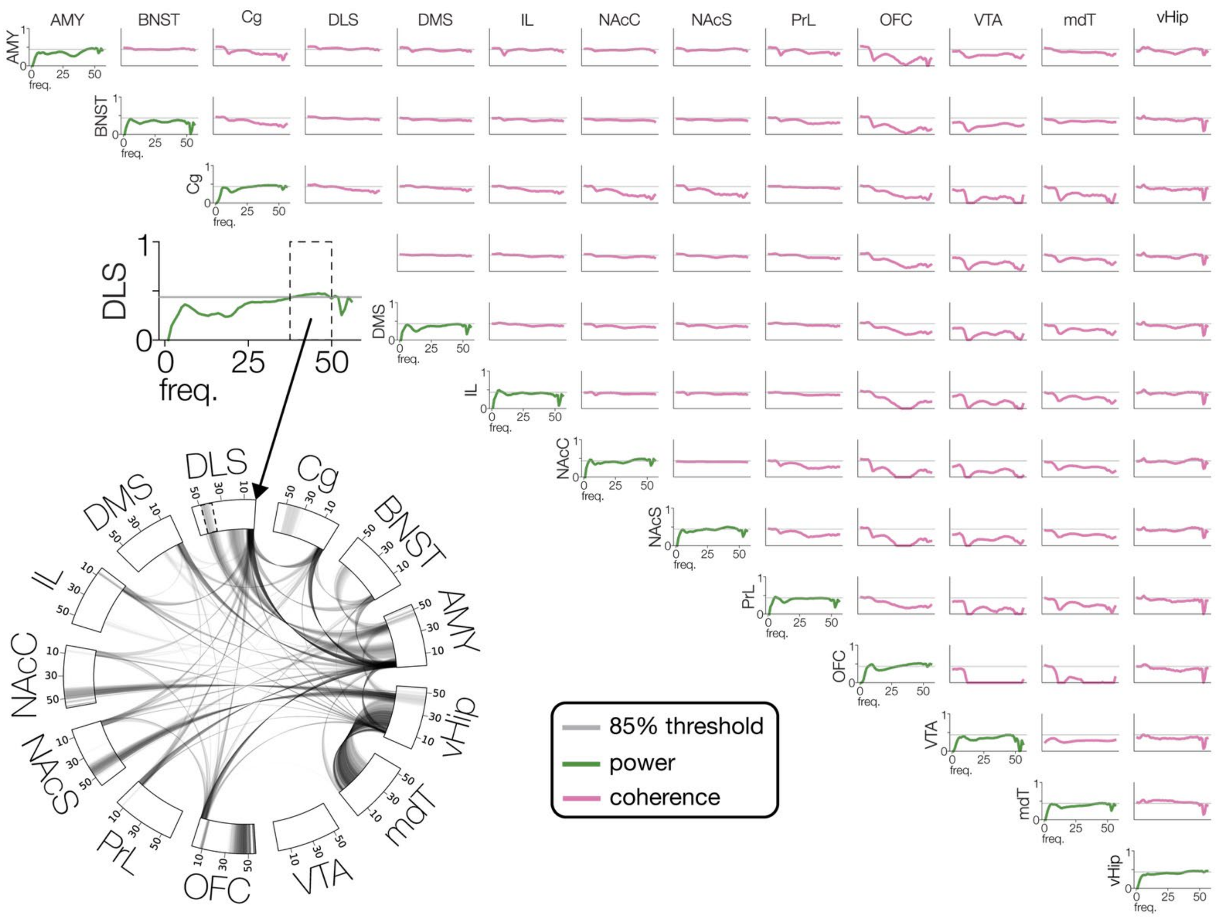


**Figure S4: Architecture of the *EN-Withdrawal* network.** The full *EN-Withdrawal* network is illustrated in a triangular plot. Diagonal plots (green) depict within-region power feature weights across frequencies, and off-diagonal plots (pink) depict inter-regional coherence feature weights. Although all features receive a weight indicating their contribution to the network, for visualization purposes we thresholded at the 85th percentile and display only the top 15% of weighted features in the Circos plot (bottom left). Spectral power features appear within the rim and coherence connections link regions across the plot. Opacity reflects feature weight magnitude. An enlarged example from the dorsolateral striatum (DLS) highlights power features exceeding the threshold between ~40–50 Hz (green line above gray cutoff), corresponding to the DLS power segment in the Circos plot (dashed boxes).

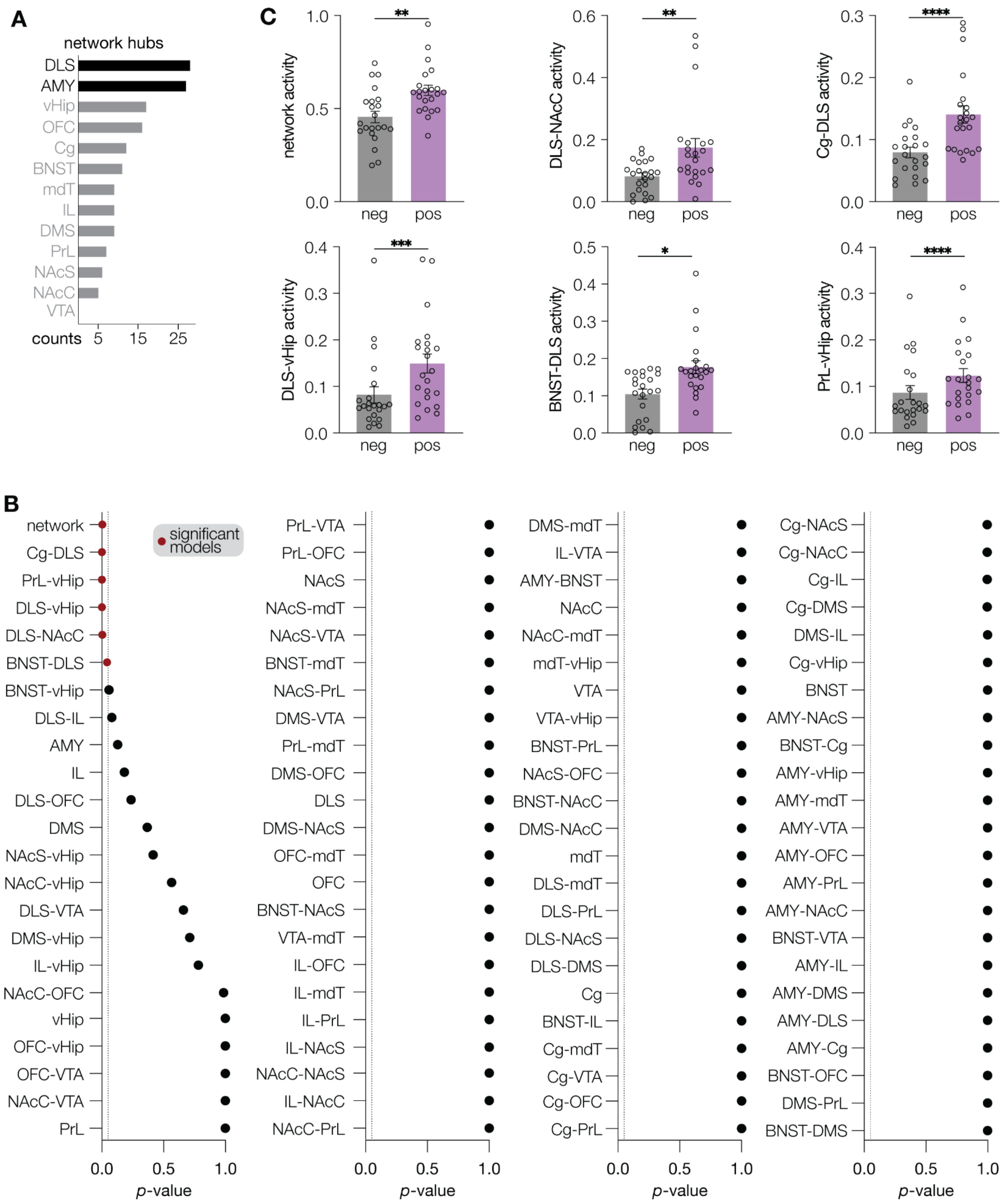


**Figure S5: Characterization of *EN-Withdrawal*.** (**A**) Network hubs, defined as the occurrence of a given brain region in the top 15% of network features, include the dorsolateral striatum and amygdala. **(B)** Bonferroni-corrected *p*-values for all single feature-group models individually trained on naloxone-precipitated fentanyl withdrawal and tested on morphine-withdrawal holdout datasets. **(C)** Average network scores per mouse for negative (non-withdrawal)- and positive (withdrawal)-class data for the six significant models from panel B ($p<0.05$; Bonferroni-corrected linear mixed-effects model). Bar graphs depict mean +/- sem. * $p<0.05$, ** $p<0.01$, *** $p<0.001$, **** $p<0.0001$.

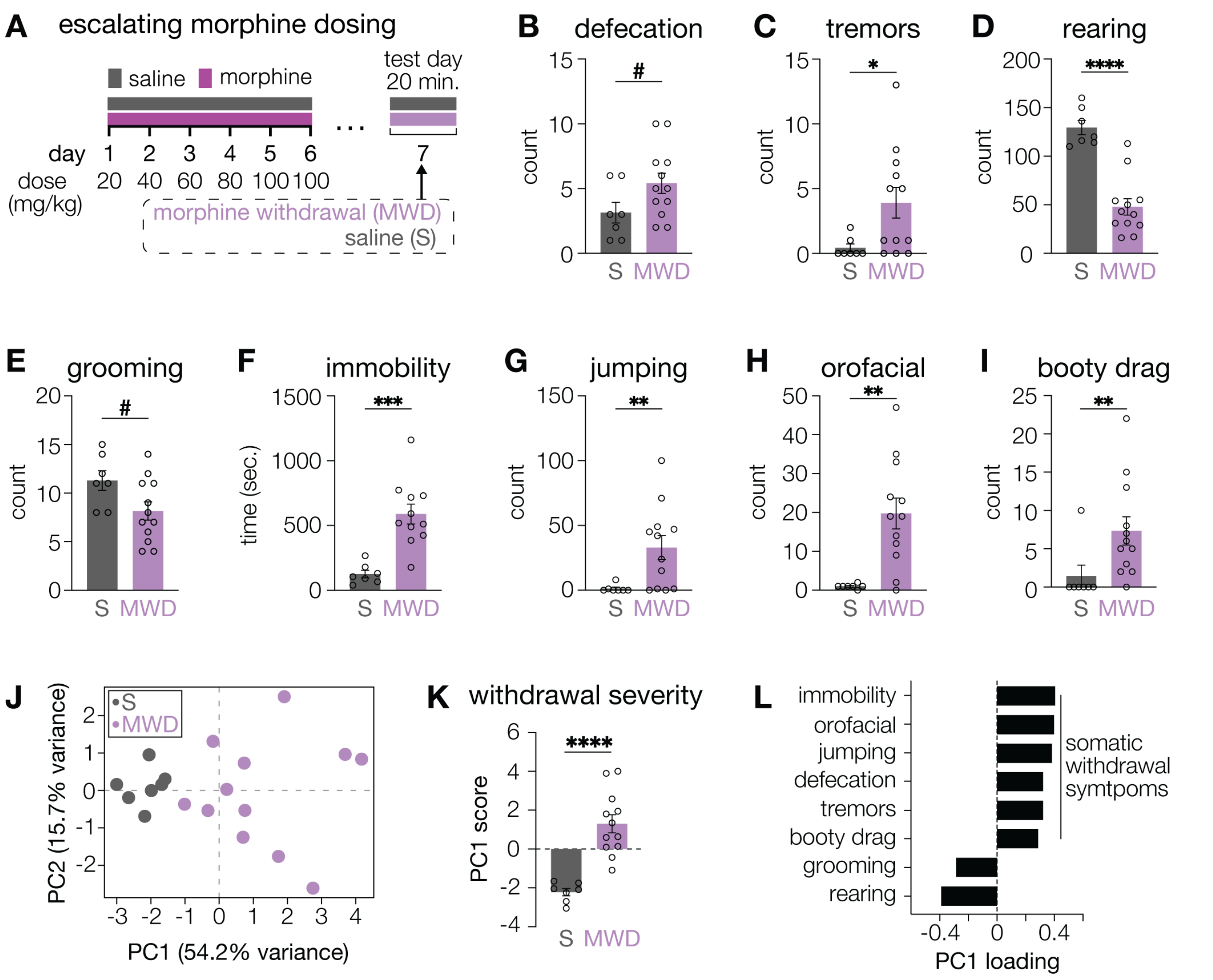


**Figure S6: Withdrawal severity index. (A)** Escalating morphine withdrawal (MWD) paradigm. Mice received twice-daily subcutaneous injections of morphine with escalating doses for 6 days, or an equal volume of saline (S). On the following day after the final injection, spontaneous withdrawal behaviors were recorded in a clear monitoring chamber. **(B–I)** Quantification of individual behaviors comparing S vs MWD: (**B**) defecation ($p$=0.059; Welch's $t$-test), (**C**) forepaw tremors ($p$=0.035; Mann–Whitney $U$), (**D**) rearing ($p$<0.0001; Welch's $t$-test), (**E**) grooming ($p$=0.052; Welch's $t$-test), (**F**) immobility ($p$=0.0003; Mann–Whitney $U$), (**G**) jumping ($p$=0.0081; Mann–Whitney $U$), (**H**) orofacial behaviors ($p$=0.002; Mann–Whitney $U$), and (**I**) booty dragging ($p$=0.0083; Mann–Whitney $U$). **(J)** Principal component analysis (PCA) of all behaviors revealed clustering of mice by treatment group. **(K)** PC1 scores stratified non-overlapping groups, with MWD mice showing significantly elevated levels of PC1 expression ($p$<0.0001; Welch's $t$-test). **(L)** PC1 loadings, hereafter referred to as the withdrawal severity index, indicate that higher scores reflect increased immobility, orofacial behaviors, jumping, defecation, forepaw tremors, and booty dragging (somatic withdrawal symptoms), and decreased grooming and rearing (healthy exploratory behaviors). Bar graphs depict mean +/- sem. # $p$<0.1, * $p$<0.05, ** $p$<0.01, *** $p$<0.001, **** $p$<0.0001

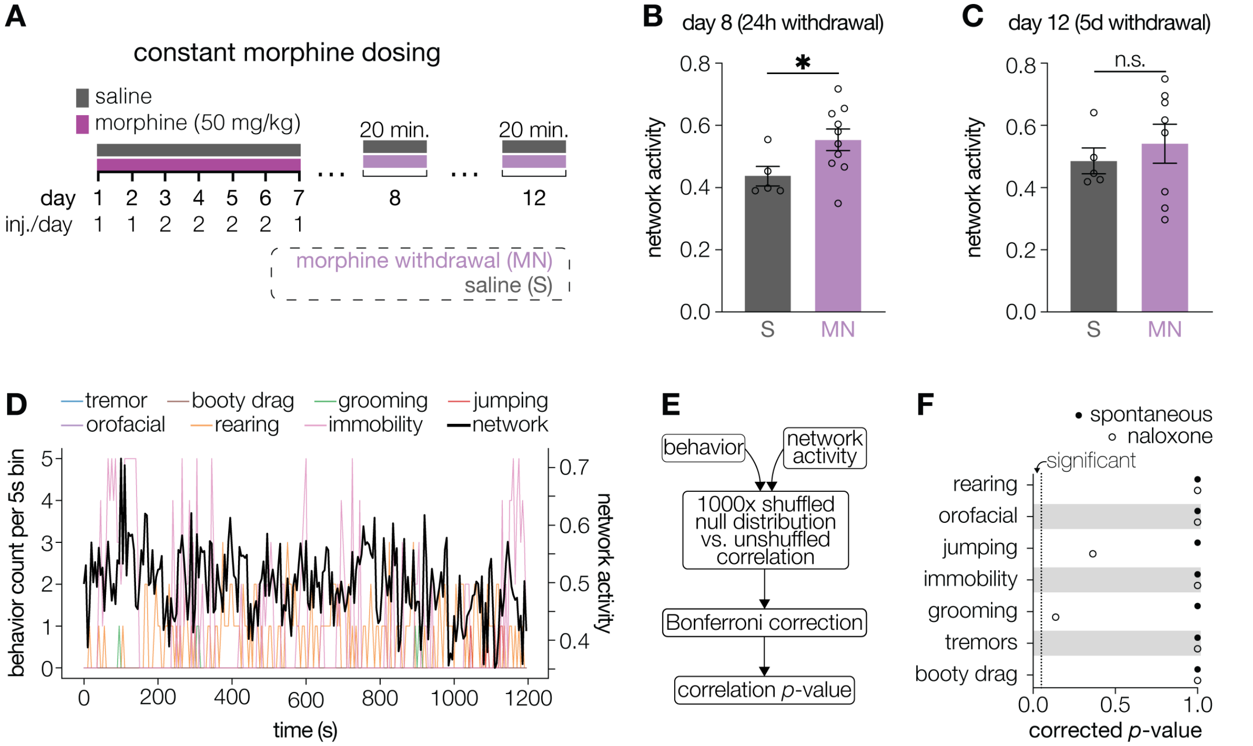


**Figure S7: Elevated *EN-Withdrawal* activity persists for at least 24 hours following naloxone-precipitated withdrawal and does not reflect ongoing behaviors. (A)** Continuation of schematic in **Figure 3A**, illustrating additional recordings 24 hours and 5 days following naloxone-precipitated withdrawal. **(B)** Network activity was significantly elevated in MN mice (N=10) compared to saline mice (N=5) 24 hours following naloxone-precipitated withdrawal ($p$=0.028; two-tailed Welch's $t$-test). **(C)** Difference in network activity between groups disappeared within 5 days of withdrawal ($N_{saline}$=5, $N_{MN}$=8; $p$=0.473; two-tailed Welch's $t$-test). **(D)** Example of behavioral and network variation in 5-second time bins. Occurrence of behavioral symptoms is plotted on the left y-axis and network activity on the right y-axis. **(E)** Statistical testing schematic. Briefly, the true correlation between any given behavior and the corresponding network activity (two-tailed Spearman correlation) is compared to a null distribution created by randomly shuffling the network activity, calculating a "shuffled" correlation, and repeating 1000 times. This is repeated for each mouse, and the average unshuffled and shuffled distributions are calculated. Then, a $p$-value is derived between these distributions, and a Bonferroni correction is applied to the final list of $p$-values to account for multiple hypothesis testing. **(F)** Bonferroni corrected $p$-values of the permutated correlation testing between any given behavior and the corresponding network activity for periods of naloxone-precipitated and spontaneous withdrawal. Dotted line indicates significance threshold at $p$=0.05. Bar graphs depict mean +/- sem. * $p$<0.05

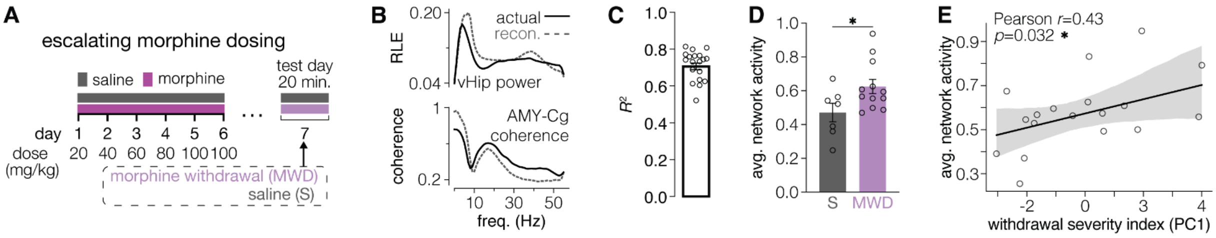


**Figure S8: *EN-Withdrawal* encodes spontaneous morphine withdrawal in two validation cohorts. (A)** Escalating morphine withdrawal paradigm – same as Figure S6. **(B)** Representative *EN-Withdrawal* feature reconstruction for ventral hippocampal power (top: relative log energy (RLE)) and amygdala–cingulate coherence (bottom). **(C)** Explained variance for reconstruction of LFP spectral features per mouse (N=19). **(D)** *EN-Withdrawal* activity is significantly elevated in withdrawing mice compared to controls ($p$=0.0219; one-tailed Welch's $t$-test). **(E)** Average *EN-Withdrawal* activity shows a positive correlation with the behavioral withdrawal severity index ($r$=0.43, $p$=0.032; one-tailed Pearson correlation). Bar graphs depict mean +/- sem. For correlation plots, the line represents the best linear fit with the shaded region indicating the 95% confidence interval. * $p$<0.05.

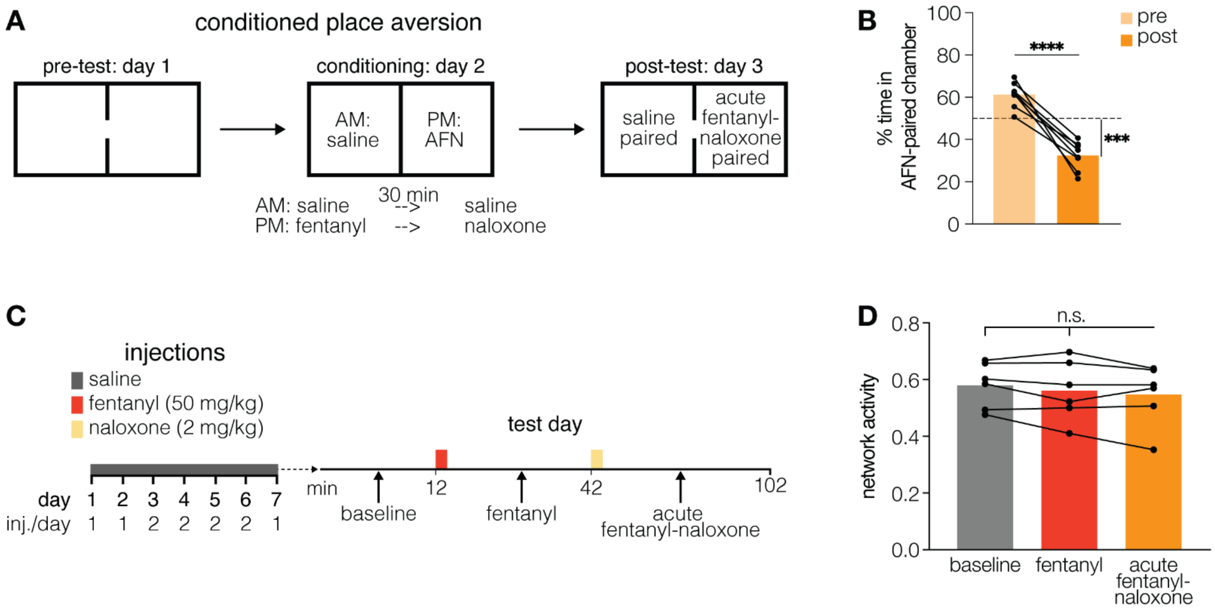


**Figure S9: *EN-Withdrawal* fails to encode aversion. (A)** Conditioned place aversion (CPA) paradigm. On day 1, mice (N=8) freely explored a two-chamber apparatus with distinct wall and floor cues to establish baseline preference. On day 2, mice received saline injections 30 min before and immediately prior to confinement in their non-preferred chamber (morning session). In the afternoon, mice received fentanyl 30 minutes prior to receiving a naloxone injection and then immediately confined to their preferred chamber (afternoon session). On day 3, mice again explored both chambers freely, and time spent in each compartment was quantified. **(B)** Mice developed a robust conditioned place aversion following a single fentanyl–naloxone pairing (N=8, $p<0.0001$; pre vs. post two-tailed paired $t$-test; $p=0.0001$; post vs. 50% chance indicated by dashed line, one-sample $t$-test). **(C)** Timeline for the electrophysiological holdout cohort (N=6) receiving acute fentanyl–naloxone dosing, mirroring the fentanyl-training paradigm in Figure 2. Mice received once- or twice-daily intraperitoneal saline injections for six days. On day 7, they were recorded for 12 min at baseline, then 30 min following a fentanyl injection, and 60 min following a naloxone injection. **(D)** *EN-Withdrawal* activity did not differ across baseline, fentanyl, or naloxone periods ($p=0.210$; repeated-measures one-way ANOVA), indicating that the network fails to encode aversive responses. Bar graphs depict mean values. *** $p<0.001$, **** $p<0.0001$.

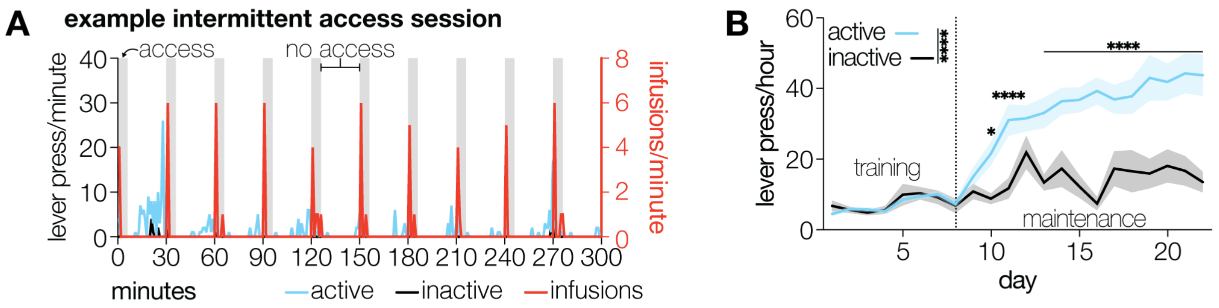


**Figure S10: Fentanyl self-administration. (A**) Example intermittent access, maintenance session from a single mouse. Here, mice undergo 10 repeating, 30-minute blocks in which animals have access to fentanyl for 6 minutes (vertical grey bars), indicated by the illumination of a cue light directly above the lever, followed by 24 minutes of no access, in which the cue light is turned off but the levers remain available. **(B)** Active and inactive lever presses across days. Notably, both levers are pressed at similar rates during training but diverge during intermittent access sessions (N=22; Day $F_{21,462}$=18.02, $p$<0.0001; Lever $F_{1,22}$=43.28, $p$<0.0001; Day x Lever $F_{21,439}$=11.29, $p$<0.0001; mixed-effects model; significant, Bonferroni-adjusted post hoc comparisons between active and inactive lever presses per day are shown on the graph.). Line graph depicts mean +/- sem. * $p$<0.05, *** $p$<0.001, **** $p$<0.0001
